# The CECR2 bromodomain displays distinct binding modes to select for acetylated histone proteins versus non-histone ligands

**DOI:** 10.1101/2024.12.09.627393

**Authors:** Margaret Phillips, Hassan Zafar, Matthew R. Marunde, Elizabeth D. Cook, Marco Tonelli, Laiba F. Khan, Amy Henrickson, James M. Lignos, Janet L. Stein, Gary S. Stein, Seth Frietze, Borries Demeler, Karen C. Glass

## Abstract

The cat eye syndrome chromosome region candidate 2 (CECR2) protein is an epigenetic regulator involved in chromatin remodeling and transcriptional control. Its bromodomain (CECR2-BRD) directs binding to acetylated histones and the acetylated RelA subunit of NF-κB, a key transcription factor in inflammatory signaling. However, the molecular basis of CECR2-BRD binding specificity, particularly with non-histone ligands, remains poorly understood. Using high-throughput peptide and nucleosome binding assays, isothermal titration calorimetry (ITC), analytical ultracentrifugation (AUC), and nuclear magnetic resonance (NMR) spectroscopy, we characterized the molecular determinants of CECR2-BRD ligand recognition. CECR2-BRD selectively binds acetylated H3 and H4 tails, with enhanced affinity for multi-acetylated targets.

Adjacent post-translational modifications (PTMs), such as phosphorylation, modulate acetyllysine recognition, particularly with histone H3 ligands. Notably, the CECR2-BRD exclusively engages nucleosomes containing multi-acetylated histones, consistent with a role in transcriptional activation. We further demonstrate that the CECR2-BRD interacts with acetylated RelA using the same conserved binding pocket used for histones, yet it shows a distinct residue perturbation pattern. The conformational flexibility observed in the CECR2-BRD indicates it uses an adaptive mechanism to distinguish between acetylated histone and non-histone substrates. Collectively, our results establish the CECR2-BRD as a key chromatin reader and a promising therapeutic target for modulating transcription and NF-κB-mediated inflammatory pathways.

**HIGHLIGHTS:** • The CECR2-BRD recognizes short-chain acyl modifications on histone H3 and H4 N-terminal tails.

• The CECR2-BRD binds acetylated RelA ligand with micromolar affinity.

• NMR perturbation studies delineate the distinct binding modes driving CECR2-BRD recognition of histone versus non-histone ligands.

• Site-directed mutagenesis reveals the specificity determinants of CECR2-BRD ligand binding.

• The CECR2-BRD strongly interacts with only multi-acetylated nucleosomes.

## INTRODUCTION

CECR2 is a regulatory subunit of the ATP-dependent chromatin remodeling complex CERF (CECR2 containing remodeling factor)[1]. Together with imitation switch (ISWI) ATPases, and the closely related sucrose non-fermenting family members SNF2L and SNF2H, CECR2 regulates nucleosome organization to facilitate DNA accessibility during neurulation, spermatogenesis, and other developmental processes[1, 2]. These chromatin remodeling complexes also play critical roles in transcription and replication by modulating nucleosome spacing[3, 4]. Additionally, CECR2 contributes to the DNA damage response through its bromodomain, which inhibits γ-H2AX activity[5].

Inflammation plays an essential role in the initiation and progression of cancer[6], and the nuclear transcription factor NF-κB is a crucial mediator of inflammatory responses in carcinogenesis[7]. The most abundant, activated form of NF-κB is composed of a heterodimer of the p65 (RelA) and p50 (NF-κB1) subunits[8]. Acetylation of RelA at lysine 310 is required for optimal transcriptional activity of NF-κB[9]. A recent study found that CECR2 interacts with acetylated RelA, enhancing NF-κB signaling and promoting the expression of pro-inflammatory and pro-metastatic factors[10]. Moreover, inhibition of CECR2-BRD binding activity suppresses NF-κB mediated immune response and pro-metastatic genes[10]. Elevated CECR2 expression has also been identified as a potential poor prognostic marker for patients with glioma and breast cancer[11, 12]. Thus, targeting CECR2 and its bromodomain activity represents a promising strategy for modulating NF-κB signaling in cancer.

Full-length CECR2 is a multidomain protein comprising an N-terminal DDT domain (DNA-binding homeobox and different transcription factors), an Adenine-Thymine-rich DNA binding motif (AT-hook), and a bromodomain[4, 13]. The DDT and AT-hook domains in CECR2 have been shown to bind DNA[14, 15], whereas its bromodomain recognizes acetylated lysine[16]. Although the CECR2-BRD is known to recognize acetylated histones, a comprehensive molecular understanding of how it ‘reads’ acetyllysine in histone and non-histone contexts is lacking. The kinetic and conformational determinants that govern CECR2-BRD’s recognition of diverse acetyllysine modifications have yet to be elucidated at high resolution. Furthermore, how neighboring combinatorial post-translational modifications (PTMs) influence CECR2-BRD binding activity and regulate its function remains largely unexplored. Over the past decade, research has demonstrated that bromodomains are excellent drug targets due to their central role in gene regulation, chromatin remodeling, and protein-protein interactions[17]. Elucidating the specificity determinants driving CECR2-BRD ligand binding interactions in these processes would provide valuable insights for the development of selective inhibitors to modulate its activity.

Here, we investigate the molecular mechanisms underlying acetyllysine recognition by the CECR2-BRD using a combination of high-throughput Captify™-Alpha peptide and nucleosome binding assays, ITC, and NMR spectroscopy. Our analyses reveal that the CECR2-BRD recognizes short-chain acylated histone H3 and H4 ligands, displays enhanced binding to multi-acetylated substrates in both peptide and nucleosome contexts, and engages RelA through the same conserved binding pocket while exhibiting ligand-specific conformational changes. Together these findings provide new insights into how the CECR2-BRD discriminates acetyllysine marks among histone and non-histone substrates.

## METHODS

### Plasmid construction

A construct encoding amino acids 424-538 covering the bromodomain (BRD) region of the human CECR2 protein (UniProt code: Q9BXF3-1) was codon optimized for expression in *Escherichia coli*, and cloned into a pGEX-6P-1 plasmid containing an N-terminal GST tag and a PreScission Protease cleavage site (GenScript, Piscataway, NJ, USA). The plasmid was transformed into *E. coli* BL21(DE3) pLysS competent cells (Novagen, MA, USA) for protein expression.

### Protein expression and purification

The GST-CECR2-BRD containing BL21(DE3) pLysS cells were cultivated following established protocols[18, 19]. Briefly, cells were grown in 2 L of Terrific broth (TB) media, or ^15^N ammonium chloride and ^13^C-glucose-supplemented minimal media, with ampicillin and chloramphenicol antibiotics added, at 37°C. Once the optical density at 600 nm (O.D. 600) reached 0.6, the temperature was reduced to 20°C. Subsequently, the cells were grown until the O.D. 600 reached a range of 0.8-1.0 and then induced with 0.1 mM isopropyl β-D-1-thiogalactopyranoside (IPTG). The induced cells were further grown overnight at 20°C. Following induction, the cells were harvested by centrifugation at 5,000 RPM for 10 minutes at 4°C. The resulting cell pellet was resuspended in 100 mL of lysis buffer, which contained 50 mM Tris−HCl pH 7.5, 500 mM NaCl, 5% glycerol, 0.05% Nonidet P-40 alternative, 1 mM dithiothreitol (DTT) and supplemented with 0.1 mg/mL of lysozyme and 1 tablet of protease inhibitor (Pierce protease inhibitor tablets, EDTA-free, ThermoFisher). The cells were then lysed using sonication, and the resulting cell lysate was centrifuged at 10,000 RPM for 20 minutes. The supernatant was mixed with glutathione agarose resin (ThermoFisher) and incubated at 4°C with gentle agitation for 1 hour. Afterward, the suspension was transferred to Econo-Column Chromatography Columns (Bio-Rad) and washed with wash buffer (20 mM Tris−HCl pH 7.5, 500 mM NaCl, 5% glycerol, and 1 mM DTT). The GST tag was then cleaved by incubating the washed beads in wash buffer supplemented with PreScission Protease (GE Healthcare) overnight at 4°C. The eluted CECR2-BRD protein was subsequently concentrated to a total volume of approximately 3 mL. For Captify-Alpha assays, the GST-tag was retained by eluting with 50 mM reduced glutathione and free glutathione removed via dialysis (48 hours) into 20 mM Tris-HCl pH 8.0, 200 mM NaCl, 1 mM DTT, and 20% glycerol. The protein concentration was determined by measuring the protein absorbance at 280 nm. The purity of the CECR2-BRD was confirmed through sodium dodecyl sulfate-polyacrylamide gel electrophoresis (SDS-PAGE) stained with GelCode Blue Safe protein stain (ThermoFisher).

### Captify-Alpha Binding Assays

The approach was performed as previously described with minor modifications[19, 20]. In brief, 5 μL GST-CECR2-BRD was combined with 5 μL histone peptides (100 nM) or nucleosomes (10 nM) and incubated 30 minutes in a 384-well plate. Binding was detected by adding a 10 μL mixture of 2.5 μg/mL AlphaLISA glutathione Acceptor (Revvity AL109M) and 10 μg/mL streptavidin Donor beads (Revvity 6760002). After a 60-minute incubation, Alpha counts were measured on a PerkinElmer 2104 Envision plate reader (680 nm laser excitation, 570 nm emission filter ± 50 nm bandwidth). Reagents for peptide assays were prepared in: 50 mM Tris pH 7.5, 50 mM NaCl, 0.01% Tween-20, 0.01% BSA, 1 mM TCEP, 0.0004% Poly-L-Lysine. For nucleosomes they were prepared in: 20 mM Tris pH 7.5, 250 mM NaCl, 0.01% NP-40, 0.01% BSA, 1 mM DTT. Binding curves were plotted in GraphPad Prism 10 using 4- parameter logistic nonlinear regression curves. Relative EC_50_ (EC_50_ ^rel^) values were computed in GraphPad Prism; the concentration of GST-CECR2-BRD necessary to elicit 50% of maximal binding signal. Assay points exceeding the hook point were excluded from EC_50_ ^rel^ calculations[20, 21]. All binding interactions were performed in duplicate, and incubations performed at room temperature under subdued lighting.

### Peptide synthesis

All modified and unmodified peptides were obtained from GenScript (Piscataway, NJ, USA). All histone H4 peptides (res 1-24) were synthesized with N-terminal acetylation (Nα-ac) and C-terminal amidation, while all H3 peptides (res 1-24) contain only a C-terminal amidation. The RelA peptides (305-315) contain no N-term acetylation or C-term amidation. All peptides were purified with HPLC to reach 98% purity and confirmed using mass spectrometry analysis. The amino acid sequences of each peptide can be found in **Table 1** and **Table 3**.

**Table 1:**
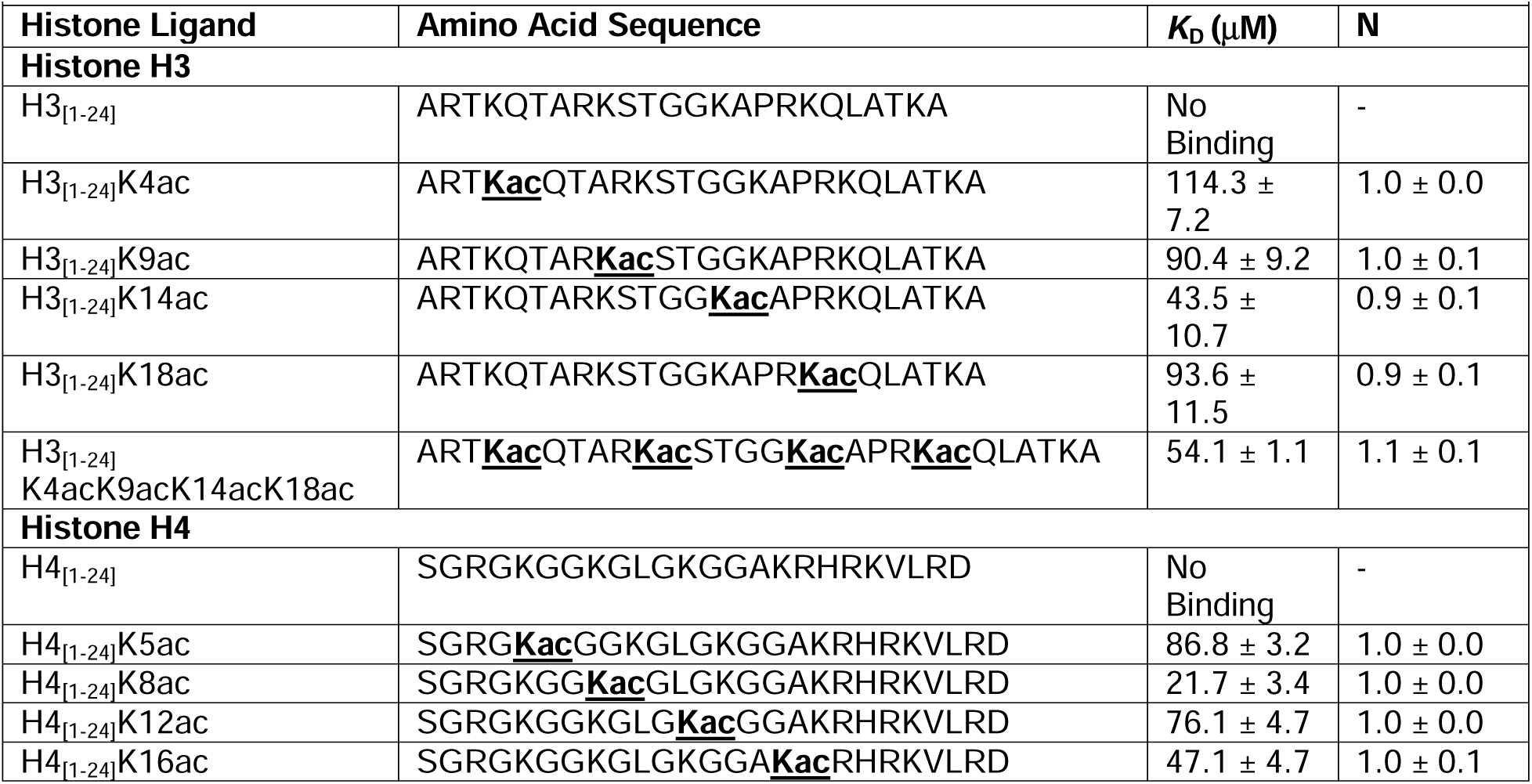
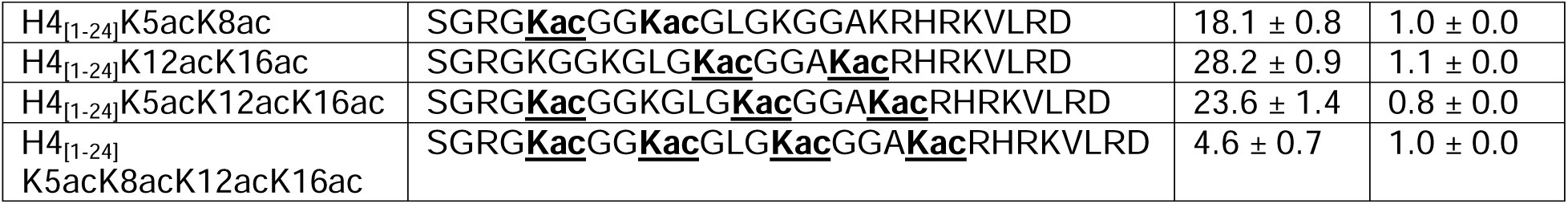
CECR2 bromodomain binding to acetylated histone H3 and H4 ligands measured by ITC. The binding affinities are represented as K_D_ (μM), and N denotes stoichiometry. All peptides include residues 1 to 24.

**Table 2:**
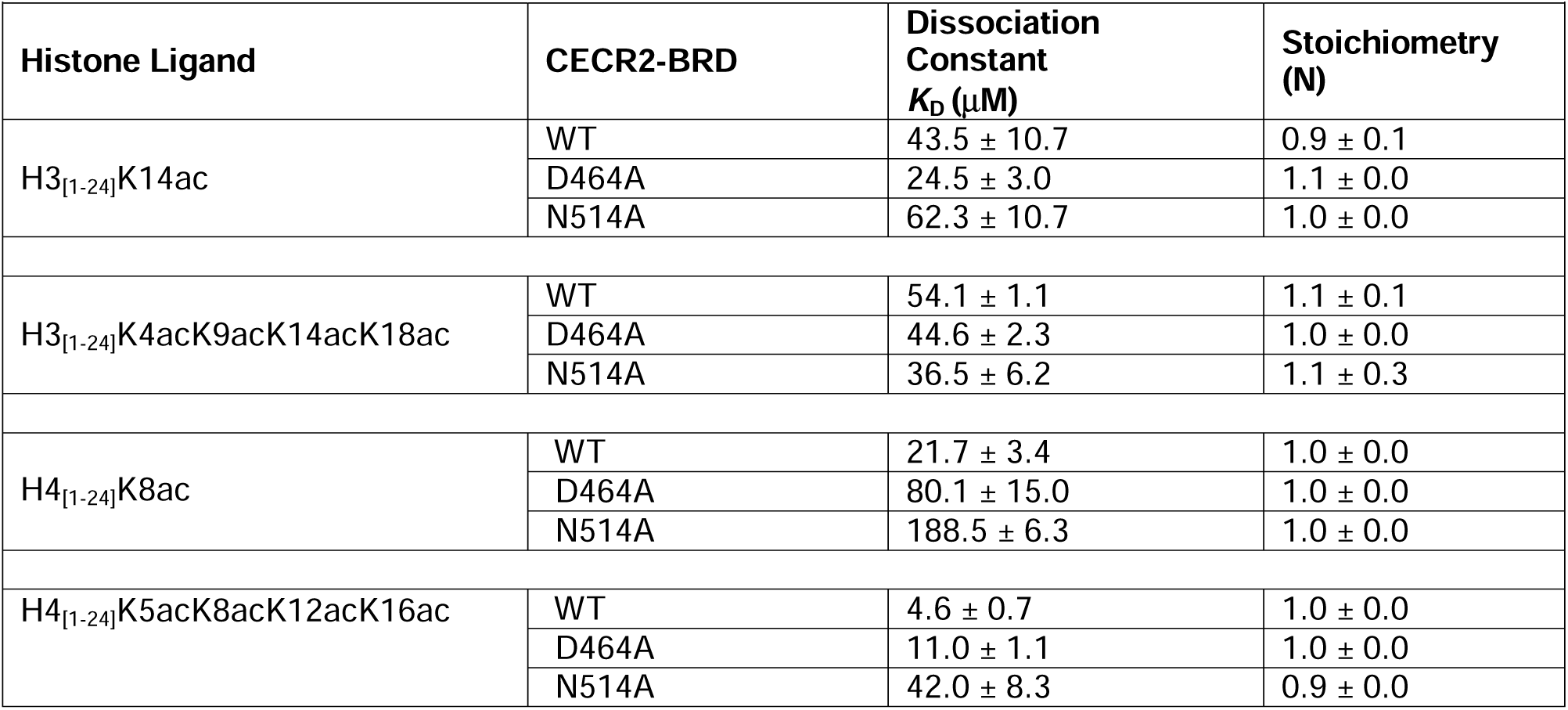
CECR2 bromodomain mutants binding to acetylated histone H3 and H4 ligands measured by ITC. The binding affinities are represented as *K*_D_ (μM), and N denotes stoichiometry. All peptides include residues 1-24.

| **by ITC.** The binding affinities are represented as $K_D$ ($\mu\text{M}$), and N denotes stoichiometry. All peptides include residues 1-24. | | | |
| --- | --- | --- | --- |
| Histone Ligand | CECR2-BRD | Dissociation Constant $K_D$ ($\mu\text{M}$) | Stoichiometry (N) |
| H3<sub>[1-24]</sub>K14ac | WT | $43.5 \pm 10.7$ | $0.9 \pm 0.1$ |
| | D464A | $24.5 \pm 3.0$ | $1.1 \pm 0.0$ |
| | N514A | $62.3 \pm 10.7$ | $1.0 \pm 0.0$ |
| H3<sub>[1-24]</sub>K4acK9acK14acK18ac | WT | $54.1 \pm 1.1$ | $1.1 \pm 0.1$ |
| | D464A | $44.6 \pm 2.3$ | $1.0 \pm 0.0$ |
| | N514A | $36.5 \pm 6.2$ | $1.1 \pm 0.3$ |
| H4<sub>[1-24]</sub>K8ac | WT | $21.7 \pm 3.4$ | $1.0 \pm 0.0$ |
| | D464A | $80.1 \pm 15.0$ | $1.0 \pm 0.0$ |
| | N514A | $188.5 \pm 6.3$ | $1.0 \pm 0.0$ |
| H4<sub>[1-24]</sub>K5acK8acK12acK16ac | WT | $4.6 \pm 0.7$ | $1.0 \pm 0.0$ |
| | D464A | $11.0 \pm 1.1$ | $1.0 \pm 0.0$ |
| | N514A | $42.0 \pm 8.3$ | $0.9 \pm 0.0$ |

**Table 3:** CECR2 bromodomain mutants binding RelA_[305-315]_K310ac measured by ITC. The binding affinities are represented as *K*_D_ (μM), and N denotes stoichiometry. The peptide includes RelA residues 305-315.

| Ligand | Amino Acid Sequence | CECR2-BRD | Dissociation Constant $K_D$ (μM) | Stoichiometry (N) |
| --- | --- | --- | --- | --- |
| RelA<sub>[305-315]</sub>K310 | TYETFKSIMKK-NH<sub>2</sub> | WT | No Binding | - |
| RelA<sub>[305-315]</sub>K310ac | TYETFKacSIMKK-NH<sub>2</sub> | WT | 60.7 ± 4.3 | 1.0 ± 0.0 |
| | | D464A | 490.3 ± 67.0 | 1.0 ± 0.0 |
|  |  | N514A | No Binding | - |

### Isothermal titration calorimetry

The CECR2-BRD was dialyzed for 48 hours into 20 mM sodium phosphate buffer, pH 7.0, 150 mM NaCl, and 1 mM tris(2-carboxyethyl) phosphine (TCEP) (ThermoFisher). All ITC experiments were performed at 5°C using a MicroCal PEAQ-ITC instrument (Malvern Analytical). All ligands were loaded into the syringe at concentrations ranging from 0.6 mM to 2.5 mM, while the CECR2-BRD was present in the sample cell at concentrations ranging from 0.05 mM to 0.10 mM. The histone ligand was injected into the sample cell in a series of 20 individual, 2 μL injections, with intervals of 150 s and continuous stirring at 750 RPM. Prior to these injections, a 0.4 μL preliminary injection of the peptide ligand was made but excluded from data integration and binding constant calculation. Control runs were performed by injecting histone peptide into the protein buffer under the same conditions, and the resulting heat of dilution was subtracted from the ligand-protein runs to correct for this effect. Data analysis was conducted using MicroCal PEAQ-ITC software. A single set of binding site models was applied to fit the data and determine the stoichiometry (N) and binding constant (*K_D_*). Binding experiments were performed in triplicate, while non-binding runs were conducted in duplicate. The mean *K*_D_ value was calculated from the average of the three binding runs, and the standard deviation was determined based on the mean. All *K*_D_ values and their corresponding stoichiometries are reported in **Table 1** and **Table 3**.

### Analytical ultracentrifugation

To determine the interaction potential between the CECR2-BRD and mono- and multi-acetylated histone peptide ligands, and their effect on CECR2-BRD oligomerization, we analyzed mixtures between the two molecules by sedimentation velocity (SV) at various molar ratios including:

1. CECR2-BRD (apo), measured at 233 nm
2. CECR2-BRD + H3_[1-24]_K4acK9acK14acK18ac at a 1:2.3 molar ratio, measured at 236 nm
3. CECR2-BRD + H3_[1-24]_K4acK9acK14acK18ac at a 1:2 molar ratio, measured at 232 nm
4. CECR2-BRD + H3_[1-24]_K4acK9acK14acK18ac at a 1:0.5 molar ratio, measured at 238 nm
5. CECR2-BRD + H3_[1-24]_K14ac at a 1:0.5 molar ratio, measured at 239 nm
6. CECR2-BRD + H4_[1-24]_K5acK8acK12acK16ac at a 1:0.5 molar ratio, measured at 232 nm
7. CECR2-BRD + H4_[1-24]_K5acK8acK12acK16ac at a 1:1 molar ratio, measured at 235 nm
8. CECR2-BRD + H4_[1-24]_K8ac at a 1:0.5 molar ratio, measured at 239 nm

The CECR2-BRD protein concentration ranged from 15.6-75.3 μM, and multiple molar ratios were used in order to ensure the ligand concentrations would be above the *K*_D_ of the protein:ligand complex as determined by ITC (**Table 1**).

All samples were measured in UV-intensity mode, and to optimize signal-to-noise ratio and maintain all signals within the dynamic range of the absorbance detector, the wavelength of data collection was optimized for optical densities between 0.5-0.8 OD determined by UV spectroscopy using a ThermoFisher Genesys 10s benchtop spectrophotometer prior to AUC analysis. All samples were measured in a buffer containing 150 mM NaCl, 20 mM NaPO_4_, adjusted to pH 7.0, with 1 mM TCEP added to prevent cysteine oxidation. The experiment was performed at 35,000 RPM for 15 hours at 5°C. Intensity data were collected in an Optima AUC at the Canadian Center for Hydrodynamics at the University of Lethbridge. All data were processed with UltraScan[22], as described previously[23]. Experimental data were fitted using the 2- dimensional spectrum analysis (2DSA)[24], in a multi-step refinement process to eliminate time- and radially-invariant noise, and to define boundary conditions. The refined data were then further analyzed using the 2DSA iterative (IT) and 2DSA Monte Carlo (MC) methods[25]. To compare the sedimentation profiles of each sample, diffusion-corrected integral sedimentation coefficients were generated and overlaid using the van Holde-Weischet method[26].

### Nuclear magnetic resonance spectroscopy

CECR2-BRD (residues 424-538) was expressed and purified for NMR experiments following previously established methods[18, 19]. Briefly, uniformly labeled ^15^N CECR2-BRD samples in buffer containing 20 mM Tris-HCl pH 6.8, 150 mM NaCl, 1 mM TCEP, and 10% D_2_O were purified at 0.30 mM for chemical-shift perturbation experiments.

All two-dimensional (2D) ^15^N-HSQC (heteronuclear single quantum coherence) experiments were acquired at 25°C using a 600 MHz Bruker AVANCE III spectrometer equipped with a z-gradient 1.7 mm TCI probe at the National Magnetic Resonance Facility at Madison (NMRFAM). The experimental setup for all 2D experiments was kept similar to our previous studies[18, 19]. Histone and non-histone ligands were added in increasing concentrations of 1:0.5, 1:1, 1:2.5, 1:5, and 1:10 molar ratios of protein to the ligand, except Histone H4_[1-24]_K5acK8acK12acK16ac for which the highest molar concentration was kept at 1:5. Chemical Shift Perturbations (CSP) from the final titration point were calculated and normalized as described by Williamson[27], using the equation:

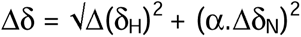

Where α = 0.2 for glycine, and 0.14 for all other residues. Δδ_H_ and Δδ_N_ represent chemical shifts in ppm for hydrogen and nitrogen atoms, respectively. CSPs were plotted as a bar graph to identify CECR2-BRD residues perturbed in response to adding specific histone ligands. These were calculated using the standard deviation (Δ ppm/ Standard deviation ppm, where standard deviation was calculated for each histone ligand) of the Euclidean chemical change of the residues as described previously[18, 19].

For the backbone assignments, ^15^N, ^13^C labeled CECR2-BRD was prepared at 0.5 mM concentration in 20 mM Tris–HCl, pH 6.8, 150 mM NaCl, 1 mM TCEP, and 10% D_2_O. 2D ^15^N-1H NHSQC, 3D HNCACB, and 3D CBCACONH experiments were recorded at 25°C on a 750 MHz Bruker AVANCE III spectrometer equipped with a z-gradient 1.7 mm TCI probe at NMRFAM. All spectra were processed using NMRPipe and analyzed using NMRFAM-Sparky and POKY[28, 29].

## RESULTS

### The CECR2 bromodomain recognizes short-chain acylated histone ligands across diverse PTM contexts

There is a limited understanding of how the CECR2 bromodomain (CECR2-BRD) interacts with modified chromatin. Using the high-throughput Captify-Alpha assay[30], we titrated GST-CECR2-BRD against H4_[1-24]_K5acK8acK12acK16ac and unmodified H4_[1-24]_ peptides, confirming a strong and selective interaction for this multiply-acetylated histone ligand (EC_50_^rel^: 7.2 nM; **Supp. Figure S1**). Using an optimized GST-CECR2- BRD probing concentration of 26 nM (**Supp. Figure S1**), we screened 287 histone peptides representing diverse sets of single and combinatorial PTMs (methylation, acylation, phosphorylation, etc.). We observed broad recognition of numerous acetylated H3, H4, H2A, and H2B ligands, with no detectable interaction to unmodified histone peptides, consistent with acetyllysine-dependent recognition (**Figure 1**). The most robust binding was observed with multi-acetylated histone H3 and H4 ligands, and the GST-CECR2-BRD tolerated adjacent methyl groups, while neighboring phosphorylation weakened interactions (**Figure 1**).

**Figure 1:**
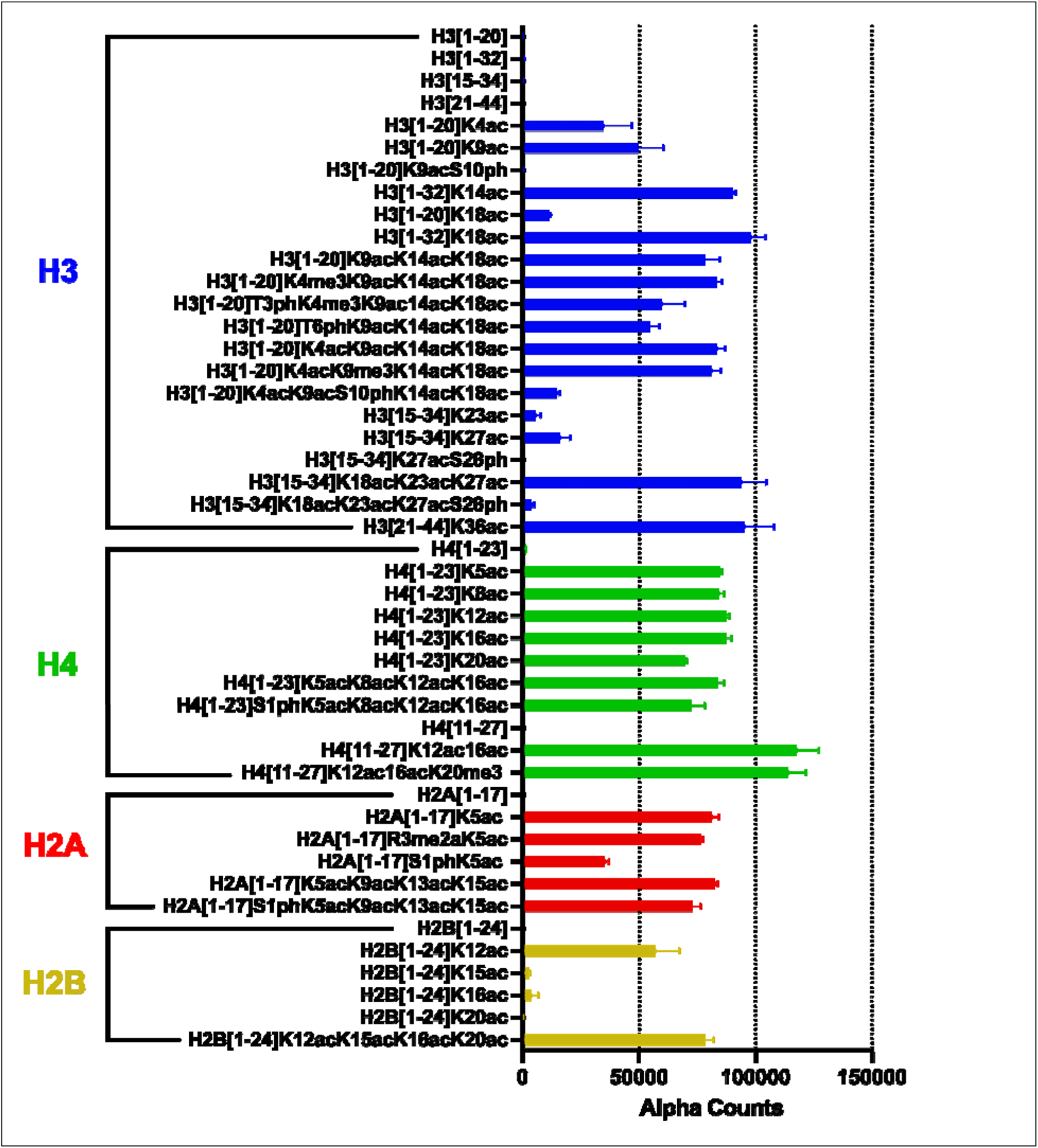
**The CECR2 bromodomain broadly recognizes acetylated histone peptides**. Using the Captify-Alpha assay, a panel of 287 histone peptides was probed with 26 nM of the GST-CECR2-BRD to assess binding interactions (refer to the ‘Captify raw data GST-CECR2- BRD.xlsx’ file for all data). The X-axis represents the Alpha Counts indicating binding strength, while the Y-axis shows the identity and amino acid coverage of each histone peptide. The GST- CECR2-BRD shows robust binding to multiply-acetylated H2A, H2B, H3, and H4 histone peptides. Binding is tolerant to adjacent methylated residues, whereas neighboring phosphorylation negatively impacts CECR2 binding to acetylated peptides. All values shown are the averages ± standard deviation (n = 2).

We next assessed whether the GST-CECR2-BRD binding pocket could accommodate bulkier acyl chains. It bound equally to acetyl, propionyl, and butyryl H3 and H4 peptides (**Figure 2**). However, interactions were weakened in the presence of crotonyl-lysine, and abrogated by bulkier acyl modifications such as beta-hydroxybutyryl, succinyl, valeryl, and hexanoyl (**Figure 2**). These results demonstrate that the GST-CECR2-BRD preferentially selects for short-chain acyl modifications.

**Figure 2.**
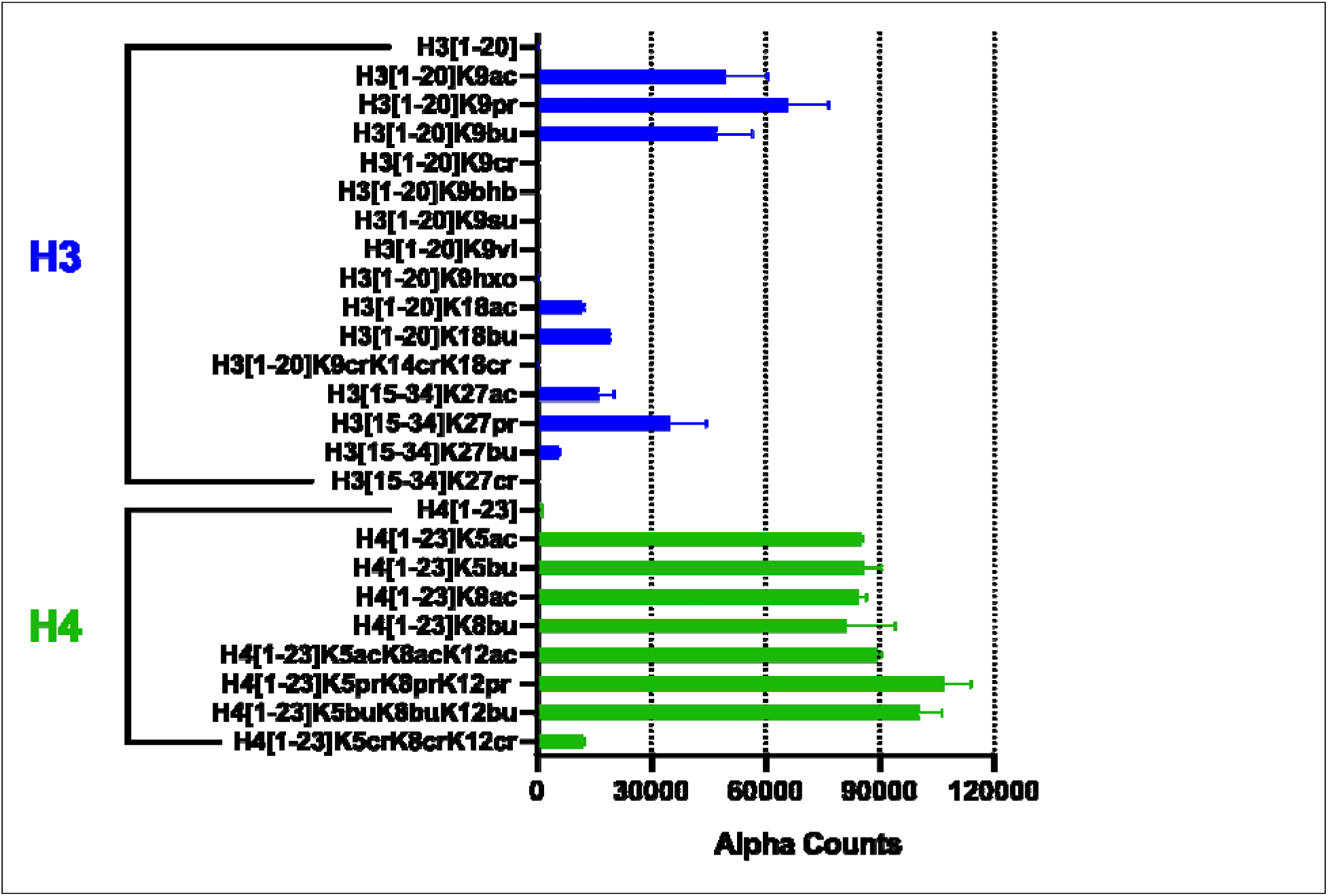
The CECR2 bromodomain recognizes acetyl, propionyl, and butyryl modifications. Binding of the GST-CECR2-BRD with longer chain acyl groups was examined using Captify-Alpha. The presence of different acyl modifications at H3K9 reveals that the GST- CECR2-BRD only interacts with acetyl (ac), propionyl (pr), and butyryl (bu) residues, as no binding was detectable with crotonyl (cr), beta-hydroxybutyryl (bhb), succinyl (su), valeryl (vl), or hexanoyl (hxo). All values shown are the averages ± standard deviation (n = 2).

While this assay is not designed to quantify peptide affinities, the CECR2-BRD displayed broad selectivity for acetylation, with limited discrimination among sequences containing multiple acetylation marks. We prioritized H3 and H4 for further biophysical studies as these peptides exhibited the highest and most reproducible signal intensity. Moreover, the H3 and H4 N-terminal tails are primary sites of combinatorial acetylation linked to transcriptional activation, and CECR2 (a component of CERF) is implicated in modulating nucleosome accessibility in acetylation-enriched chromatin contexts[1, 4, 10]. Thus, H3 and H4 provided the most biologically relevant and technically tractable targets for detailed mechanistic analysis.

### The CECR2 bromodomain prefers multi-acetylated histone H4 tails

Given the GST-CECR2-BRD’s broad recognition for acylated lysine in our initial screen (**Figure 1**), we next sought to measure binding affinities to quantitatively compare targets and identify preferred histone ligands. Using ITC, we examined the interactions of the CECR2-BRD (without a GST tag) with mono- and multi-acetylated H3 and H4 ligands (**Table 1, Supp. Figures S2 and S3**). No measurable binding was observed for the unmodified H3 and H4 peptides, confirming that acetylation is required for CECR2- BRD recognition. Among the mono-acetylated H3 peptides, CECR2-BRD exhibited relatively weak binding, with affinities ranging from *K*_D_s of 43.5 ± 10.7–114.3 ± 7.2 µM. The tetra-acetylated H3 peptide, H3_[1–24]_K4acK9acK14acK18ac, bound with a *K*_D_ of 54.1 ± 1.1 µM (**Table 1**) and therefore did not show a substantial enhancement in affinity compared with the strongest mono-acetylated H3 ligand. These results suggest that increasing the number of acetylated lysines within the H3 tail does not by itself promote substantially stronger CECR2-BRD binding.

In contrast, CECR2-BRD displayed a preference for specific acetylated H4 peptides. Among the mono-acetylated H4 ligands, H4K8ac showed the strongest interaction (*K*_D_ = 21.7 ± 3.4 µM), whereas H4K5ac, H4K12ac, and H4K16ac bound more weakly, with *K*_D_s ranging from 47.1-86.8 µM. Multi-acetylated H4 peptides generally exhibited stronger affinities, with *K*_D_ values of 18.1 ± 0.8 µM for H4K5acK8ac, 28.2 ± 0.9 µM for H4K12acK16ac, and 23.6 ± 1.4 µM for H4K5acK12acK16ac. Interestingly, the tetra-acetylated H4 peptide H4_[1–24]_K5acK8acK12acK16ac exhibited the highest affinity of all peptides tested (*K*_D_ = 4.6 ± 0.7 µM), representing an approximately five-fold increase in affinity relative to the strongest mono-acetylated H4 ligand, H4K8ac. The ITC data for CECR2-BRD reveal that it distinguishes between acetylation patterns on histone H3 and H4 tails. Whereas increased acetylation of H3 provides little additional binding advantage, extensive acetylation of the H4 tail markedly enhances CECR2-BRD recognition.

### The CECR2 bromodomain remains monomeric upon histone ligand binding

One possibility for enhanced binding to multiply-acetylated ligands is ligand-induced oligomerization of the CECR2-BRD. To test this, we performed sedimentation velocity analytical ultracentrifugation (SV-AUC) experiments to assess the oligomeric state of CECR2-BRD alone (apo control) and in the presence of histone peptides. The sedimentation coefficients of the CECR2-BRD in complex with mono- and tetra-acetylated histone ligands, including H3_[1-24]_K14ac, H4_[1-24]_K8ac, H3_[1-24]_K4acK9acK14acK18ac, and H4[1–24]K5acK8acK12acK16ac, were compared to the apo CECR2-BRD (**Figure 3**). Importantly, samples were assembled at protein:ligand ratios exceeding the ITC-derived affinities to favor complex formation. All peptide-bound sedimentation profiles closely overlapped with the apo control (**Figure 3**), indicating that the CECR2-BRD remains monomeric upon ligand binding, with no evidence of greater complex formation. Although SV-AUC cannot directly confirm peptide binding in this molecular weight range, the absence of oligomeric species rules out ligand-induced dimerization or aggregation. Together with ITC data, these results support a model where CECR2-BRD functions as a monomeric acetyllysine reader in solution.

**Figure 3:**
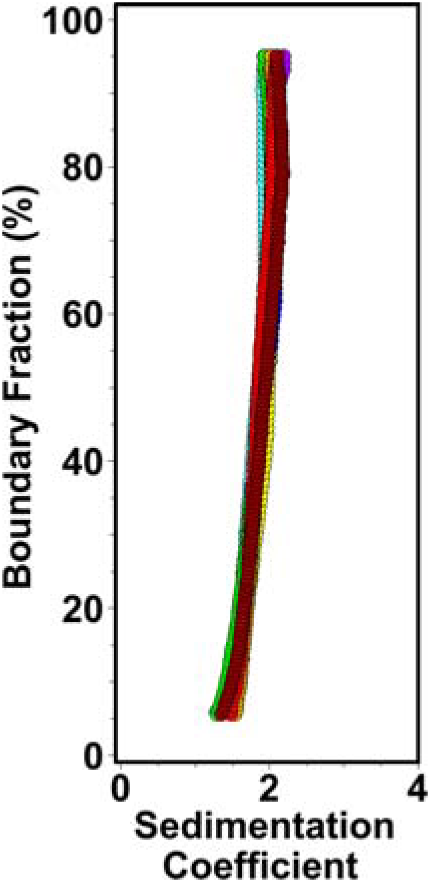
The CECR2-BRD is a monomeric acetyllysine reader in solution. Sedimentation velocity analysis of the CECR2 bromodomain alone and in complex with acetylated histone H3 and H4 ligands. The sedimentation velocity distribution van Holde-Weischet overlay plot includes: CECR2-BRD apo control (purple), CECR2-BRD + H3_[1-24]_K4acK9acK14acK18ac 1:2.3 (blue), CECR2-BRD + H3_[1-24]_K4acK9acK14acK18ac 1:2 (cyan), CECR2-BRD + H3_[1-24]_K4acK9acK14acK18ac 1:0.5 (orange), CECR2-BRD + H3_[1-24]_K14ac 1:0.5 (red), CECR2-BRD + H4_[1-24]_K5acK8acK12acK16ac 1:0.5 (green), CECR2-BRD + H4_[1-24]_K5acK8acK12acK16ac 1:1 (yellow), CECR2-BRD + H4_[1-24]_K8ac 1:0.5 (brown).

### The CECR2 bromodomain preferentially interacts with nucleosomes containing multiply-acetylated H3 and H4 histones

To evaluate CECR2-BRD binding in a more physiologically relevant chromatin context, we used the Captify-Alpha assay to assess its interaction with nucleosomes. Optimal binding conditions were first established (**Supp. Figure S4)**, after which titration of the GST-tagged CECR2-BRD revealed robust binding to ([H3.1K4acK9acK14acK18ac]_2_) and ([H4K5acK8acK12acK16ac]_2_) nucleosomes, with an EC_50_^rel^ of 6.0 nM and 9.2 nM, respectively (**Figure 4**). Thus, the GST-CECR2-BRD demonstrates a nearly identical relative interaction with tetra-acetylated H3 and H4 histones when they are in the intact nucleosome. In contrast, despite strong binding to H2A_[1-17]_K5acK9acK13acK15ac peptides (**Figure 1**), the GST-CECR2-BRD showed no detectable interaction with ([H2AK5acK9acK13acK15ac]_2_) nucleosomes. Further, no binding was observed for any nucleosomes containing histones with a single acetylation modification, including ([H4K8ac]_2_), even though the corresponding peptide exhibited a strong binding affinity (*K*_D_ = 21.7 µM, **Table 1**) among singly acetylated peptides. The pronounced binding preference for tetra-acetylated nucleosomes suggests that acetyllysine density within the nucleosomal context, rather than individual acetyllysine marks, is a critical determinant for GST-CECR2-BRD recruitment. This could reflect changes in histone tail accessibility, as the histone–DNA contacts are weakened by the charge neutralization of lysine acetylation[30–32]. These findings support a model in which the CECR2-BRD functions as a chromatin reader that preferentially engages nucleosomes marked by multiple acetylated lysines on H3 and H4, consistent with targeting transcriptionally active regions of the genome.

**Figure 4:**
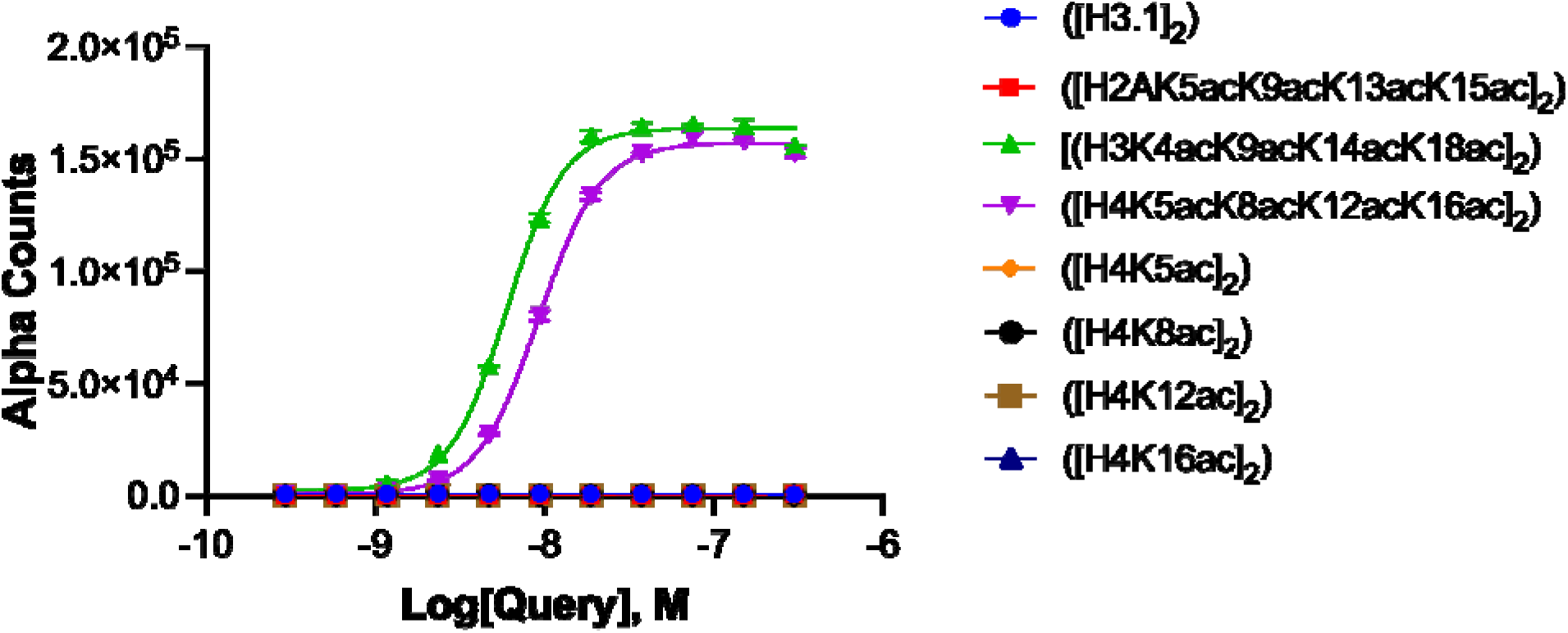
CECR2 bromodomain robustly binds multi-acetylated H3 and H4 nucleosomes. Using the Captify assay, CECR2 was titrated against nucleosomes. The X-axis represents the concentration of GST-CECR2-BRD, while the Y-axis measures the Alpha Counts indicating binding strength. The key identifies each nucleosome. GST-CECR2-BRD was assessed against single and multi-acetylated nucleosomes showing strong binding only to multi-acetylated H3 and H4 nucleosomes. All values shown are the averages ± standard deviation (n = 2).

### Molecular basis of multi-acetylated histone recognition by the CECR2-BRD

While nucleosomes represent the most physiologically relevant substrates for CECR2-BRD binding, the complexity and size of these assemblies currently limit their use in our high-resolution solution NMR analyses. Therefore, to delineate molecular contacts within the CECR2-BRD (residues 424-538) binding pocket to multi-acetylated histones, we used multi-acetylated histone H3 and H4 peptides as a simplified and structurally accessible mimic. Backbone assignments of the CECR2-BRD were completed using standard triple-resonance NMR (Methods, experiments (**Supp. Figure S5**)). Next, we performed titrations of unlabeled H3_[1-24]_K4acK9acK14acK18ac and H4_[1-24]_K5acK8acK12acK16ac peptides with ^15^N-labeled CECR2-BRD, and monitored the CSPs of CECR2-BRD residues via ^1^H-^15^N HSQC spectra (**Figure 5; Supp. Figures S6–S7**). Addition of H3_[1-24]_K4acK9acK14acK18ac or H4_[1-24]_K5acK8acK12acK16ac peptides induced comparable and significant CSPs localized to the canonical acetyllysine-binding site. Perturbations were primarily observed in residues within the ZA (res 454-479) and BC loops (res 514-518), and the adjacent αB (res 495-513) and αC helices (res 519-536). These include residues W457 and F459 of the WPF shelf, D464 in the ZA loop, and N514 in the BC loop (**Figure 5A-B**). The dramatic chemical shift observed for D464 likely arises from the aromatic ring current effect of neighboring Y467 in response to ligand binding. While the regions forming the canonical acetyllysine-binding groove are conserved across bromodomains, our CSP data validate CECR2-BRD recognition of multiply acetylated H3 and H4 ligands, define the direct contact surface for these ligands in the CECR2-BRD binding pocket, and outline the binding specificities driving ligand coordination.

**Figure 5:**
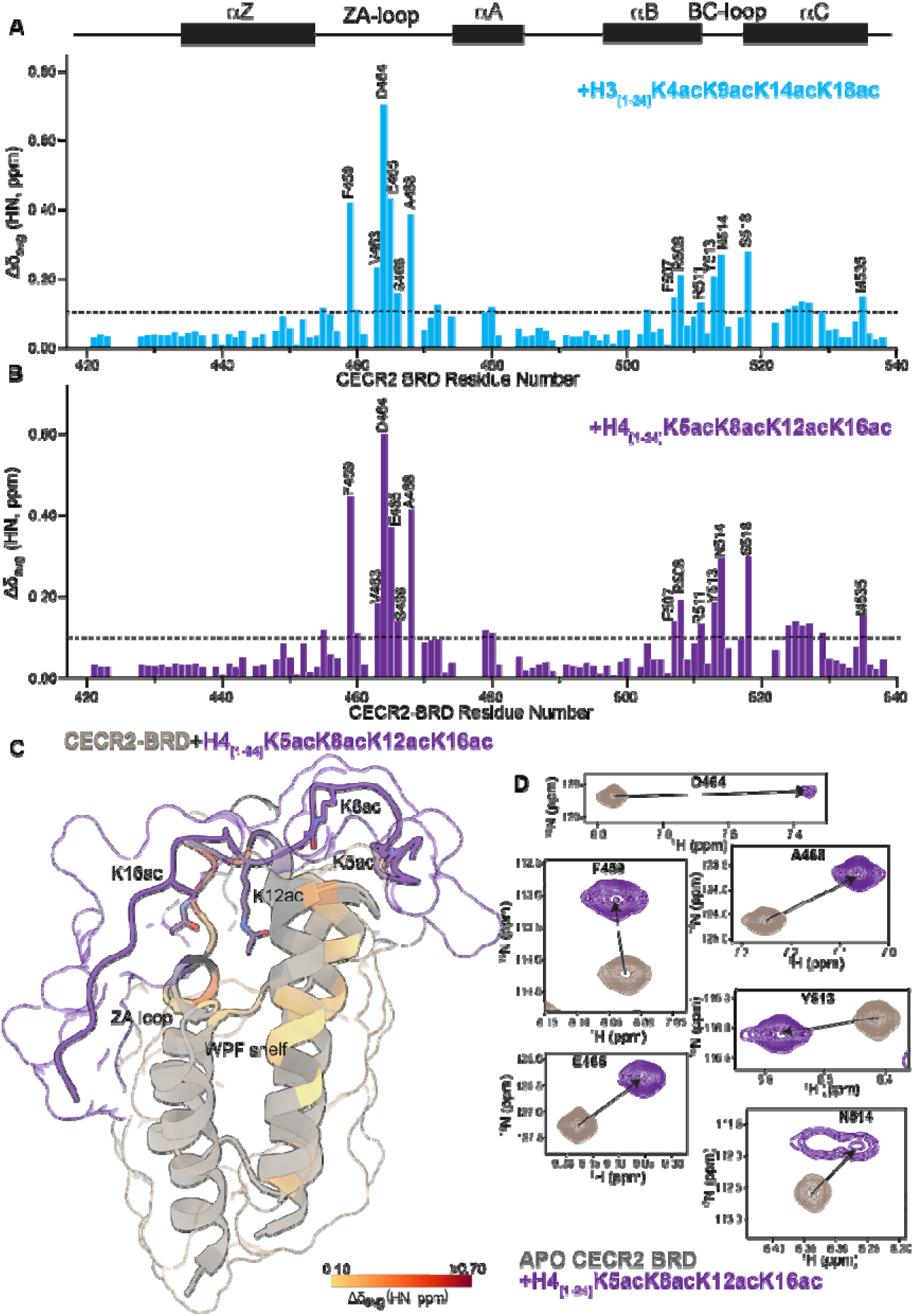
NMR characterization of multi-acetylated histone H3/H4 interaction with the CECR2 bromodomain. (A) NMR chemical shift perturbation (CSP) analysis of ^15^N-labeled CECR2 bromodomain upon titration with H3_[1-24]_K4acK9acK14acK18ac peptide at 1:10 molar ratio respectively. CSP values are plotted against CECR2-BRD residue number, highlighting strong perturbations in the ZA loop, BC loop, and WPF shelf-adjacent regions. Secondary structures within the BRD are highlighted above. (B) CSP analysis for H4_[1-24]_K5acK8acK12acK16ac at 1:10 molar ratio concentration respectively. Perturbation patterns reveal similar engagement between H3 and H4 acetyllysine with the CECR2 binding pocket residues. (C) NMR-derived chemical shift perturbations observed in H4_[1-24]_K5acK8acK12acK16ac peptide titrations are mapped onto the crystal structure of the CECR2- BRD (PDBID: 3NXB[16]), and AlphaFold3 was used to model the protein-ligand complex[33]. The bound H4 peptide (purple) is shown with acetylated lysine labeled. Regions of the CECR2- BRD experiencing the largest CSPs are colored from yellow (minimal perturbation of 0.10 ppm) to dark red (largest perturbation of ≥0.70 ppm). The ZA loop and WPF shelf are also indicated. (D) Superimposed ^1^H-^15^N HSQC spectra of select CECR2-BRD residues in the absence (beige) and presence (purple) of H4_[1-24]_K5acK8acK12acK16ac (at 1:10 molar ratio respectively) showing peak shifts. Prolines are shown in black, and unassigned residues in dark gray.

To further probe the functional relevance of key residues identified via our CSP analyses, we introduced point mutations at D464 (ZA loop, showing largest CSP) and the conserved N514 (BC loop), and compared their binding efficiency with the wild-type CECR2-BRD (**Table 2; Supp. Figure S8**). Interestingly, the D464A mutation enhances binding to H3K14ac by approximately 1.8-fold relative to WT, and slightly increases the binding affinity with H3K4acK9acK14acK18ac. In contrast, D464A reduced binding affinity for H4K8ac and tetra-acetylated H4K5acK8acK12acK16ac by approximately 3.7- and 2.4-fold, respectively, suggesting a ligand-dependent role for the D464 residue in histone H4 tail recognition. While the N514A mutation did not significantly alter the CECR2-BRD interaction with H3K14ac, it modestly enhances the affinity for tetra-acetylated H3 by approximately 1.5-fold relative to the wild-type bromodomain. In contrast, N514A substantially weakens the binding affinity for the H4K8ac and tetra-acetylated H4K5acK8acK12acK16ac ligands by 8.7- and 9.1-fold, respectively. This suggests that D464 and N514 contribute differentially to histone-tail recognition, with N514 playing a particularly important role in the recognition of acetylated H4 ligands. Together, these findings demonstrate that CECR2-BRD uses a conserved bromodomain interface to recognize acetylated H3 and H4 peptides, and its ligand binding preferences are shaped by both the number of acetyllysine modifications and histone-specific contacts. While peptide-based analyses do not fully recapitulate the 3D aspects of nucleosome binding behavior, our data provide essential insights into residue-level interactions that underlie CECR2-BRD binding specificity.

### The CECR2 bromodomain recognizes acetylated RelA via the conserved acetyllysine binding pocket

Beyond acetylated histone recognition, the CECR2-BRD has also been implicated in reading acetylated non-histone proteins, notably the transcription factor RelA (p65), which is a subunit of NF-κB[10]. Acetylation at lysine 310 (K310) is essential for RelA’s transcriptional activity and promotes the interaction of CECR2, which upregulates the NF-κB signaling pathway and enhances inflammatory gene expression in metastatic breast cancer models[9, 10]. To test whether the CECR2-BRD engages acetylated RelA similarly to acetylated histones, we first performed ITC with RelA (305– 315) peptides, with or without K310 acetylation. CECR2-BRD selectively bound the acetylated form (RelA_[305-315]_K310ac) with a moderate affinity of *K*_D_ = 60.7 ± 4.3 µM (**Table 3, Supp. Figure S9**). We also examined the effect of mutations in the CECR2- BRD binding pocket. The D464A and conserved asparagine (N514A) mutations drastically reduced and abolished binding of the CECR2-BRD with the RelA_[305-315]_K310ac ligand, respectively (**Table 3**).

The ^15^N,^1^H-HSQC NMR titration with RelA_[305-315]_K310ac shows CSPs in the canonical acetyllysine binding pocket of the CECR2-BRD, with some residues overlapping with the H3 and H4 histone binding site (**Figure 6, Supp. Figure S10**). The CSPs of residues in the BC loop region, including the conserved N514, are consistent between acetylated RelA and the acetylated histone ligands. However, we observed some subtle differences with residues in the ZA loop, where residues A468 and E465 exhibit more pronounced perturbations upon RelA_[305-315]_K310ac binding. Additionally, the CSP observed in D464 upon histone ligand binding was not seen with RelA, which instead caused a large CSP in residue N470. These differences in the CSP patterns observed in the ZA loop region of the CECR2-BRD could reflect differences in ligand-specific flexibility or dynamics between acetylated RelA and histone ligands (**Figures 5** **& 6).** Notably, our data do not imply that CECR2-BRD recognizes acetylated RelA and histones through mutually exclusive mechanisms. Rather, both interactions converge on the conserved acetyllysine recognition interface, which is likely repurposed in different cellular contexts, e.g., chromatin for histones, and transcriptional signaling for RelA. However, acetylated-RelA binding induces a unique ZA-loop perturbation pattern in the CECR2-BRD, representing a ligand-specific variation within this shared recognition framework.

**Figure 6:**
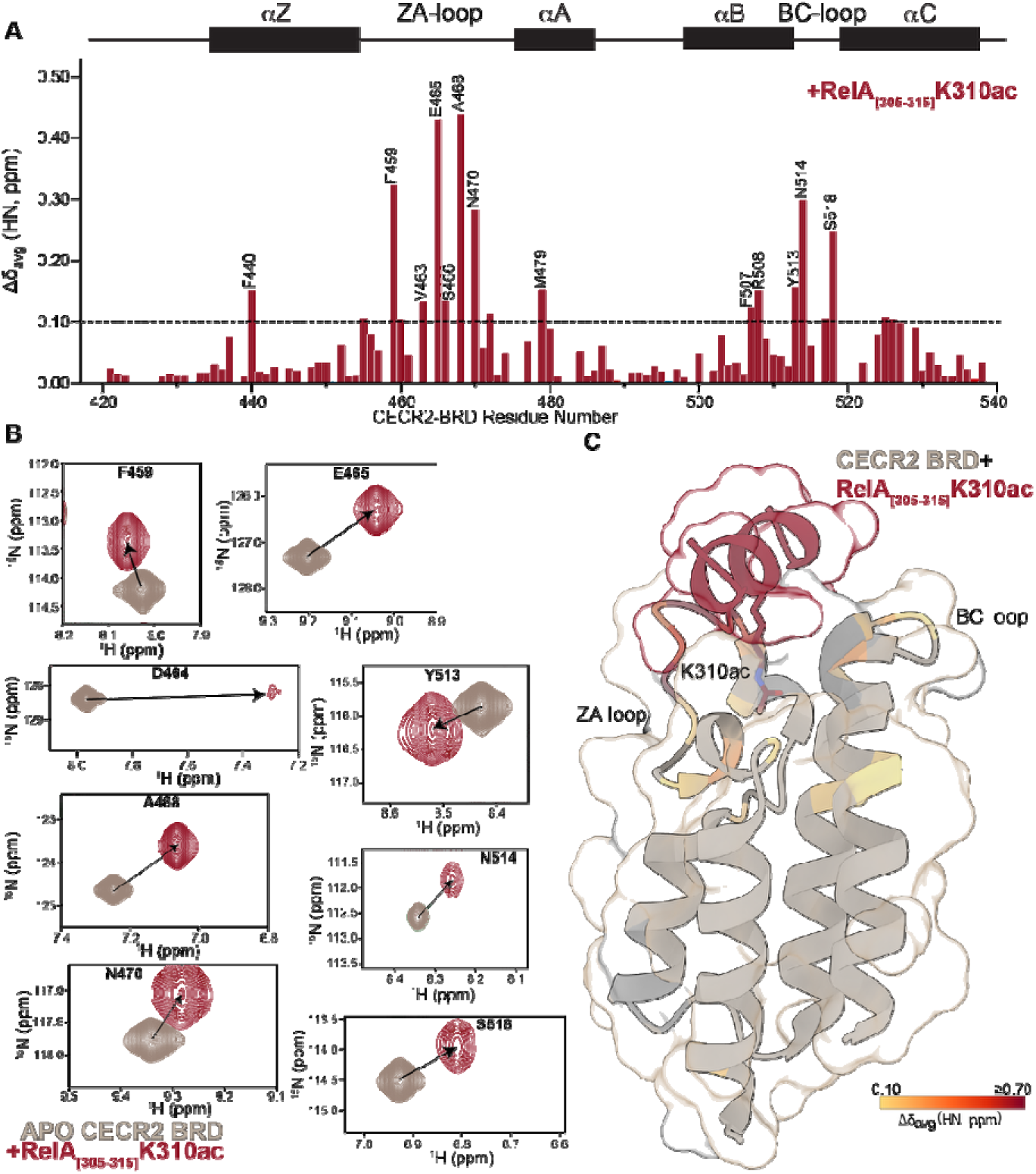
NMR characterization of RelA acetyllysine recognition by the CECR2 bromodomain. (A) NMR chemical shift perturbation (CSP) analysis of ^15^N-labeled CECR2 bromodomain upon titration with RelA_[305-315]_K310ac peptide at a molar concentration of 1:10. CSP values are plotted as a function of CECR2-BRD residue number, showing largest perturbations in the BC loop and regions adjacent to the WPF shelf. Secondary structures within the BRD are highlighted above. (B) Selected residues highlighted with overlay of ^1^H-^15^N HSQC spectra of CECR2-BRD in the absence (beige) and presence (red) of RelA_[305-315]_K310ac peptide (at a 1:10 molar ratio respectively). (C) NMR-derived chemical shift perturbations observed in RelA_[305-315]_K310ac peptide titrations are mapped onto the crystal structure of the CECR2-BRD (PDBID: 3NXB[16]), and AlphaFold3 was used to model the protein ligand complex[33]. The RelAK310ac peptide is shown in red, positioned at the canonical acetyllysine binding site. CSP magnitudes are color-coded from yellow (minimal perturbation of 0.10 ppm) to dark red (largest perturbation of ≥0.7 ppm), with BC loop, ZA loop, and WPF shelf regions indicated. Prolines are shown in black, and unassigned residues in dark gray.

## DISCUSSION

The CECR2-BRD is a key epigenetic reader that is found in the CECR2- containing remodeling factor (CERF), where it works in conjunction with the SNF2L ATP-dependent chromatin remodeler, and contributes to transcriptional regulation and DNA replication[1, 34]. However, the molecular basis by which CECR2 binds and distinguishes histone and non-histone ligands has remained unclear. In this study, we used biochemical and biophysical approaches to systematically dissect the ligand-binding preferences and structural determinants of CECR2-BRD. Our findings reveal that the CECR2-BRD preferentially binds multi-acetylated histone H3 and H4 tails and engages acetylated RelA through a conserved but adaptable binding pocket. These insights elucidate how CECR2-BRD discriminates between acetyllysine marks in chromatin and transcription factor substrates, and positions CECR2 as a potential therapeutic target for modulating gene expression and NF-κB-mediated inflammatory signaling.

We profiled CECR2-BRD binding activity using a library of histone peptides to reveal its preference for short-chain acyl modifications on histones H3 and H4, including acetyl, propionyl, and butyryl, while the bromodomain binding was weakened or abolished towards bulkier acyl chains (**Figures 1** **&** 2). This finding agrees with prior peptide data showing the CECR2-BRD recognizes butyryl and propionyl, but not crotonyl modifications on H4[35]. Additionally, histone modifications adjacent to the acetyllysine sites further modulated binding as phosphorylation hindered recognition, while methylation had little impact. This suggests that phosphorylation potentially acts as a regulatory switch to suppress CECR2 recruitment. Histone H3 phosphorylation is associated with chromatin compaction and cell cycle progression, generating a chromatin state that is less accessible to transcription factors and chromatin binding/remodeling proteins such as CECR2[36]. Reduced CECR2-BRD binding in the presence of phosphorylated H3 may also reflect a broader physiological role beyond cell-cycle control. Garcia-Pichardo *et al.* and Sharma *et al.* found that phosphorylation of histone H3 leads to impaired DNA repair[37, 38], and CECR2 is known to play a role in the DNA damage response[5]. Together, these data support a model in which the CECR2-BRD contributes to nuanced regulatory control by selectively recognizing short-chain acylation and sensing the context created by neighboring PTMs. Moreover, the position of acetyllysine marks on the histone tail modulates the CECR2-BRD binding affinity, as demonstrated by the preferential recognition of H3K14ac and H4K8ac ligands over other single acetyllysine modifications on histone H3 and H4. Similarly, oncohistone mutations that alter residues surrounding the acetyllysine sites can perturb bromodomain recognition [39, 40].

Notably, acetylation of histone H3 and H4 has been linked to neural differentiation[41, 42], and recognition of acetylated H3 histone tails by the CECR2-BRD may contribute to its recruitment to transcriptionally active chromatin to support CECR2’s role in neurulation[1]. This possibility is supported by our ITC (**Table 1**) and nucleosome-binding assays (**Figure 4**), which demonstrate recognition of multi-acetylated H3 and H4 histones. Hyperacetylation of histone H3 and histone H4 typically occurs in euchromatic regions where genes are actively expressed[43]. Histone H3 acetylation is directly associated with transcriptional activation through interaction with transcription factors, while histone H4 acetylation is indirectly associated with transcriptional activation by impacting the structure and stability of the nucleosome[31, 44, 45]. Our ITC and nucleosome binding data demonstrate that the CECR2-BRD binds both tetra-acetylated H3 and H4 peptides and recombinant nucleosomes. However, the isolated bromodomain shows a clear preference for the multi-acetylated H4 tail. It binds the tetra-acetylated H4 peptide ∼12-fold more tightly than the tetra-acetylated H3 peptide, whereas tetra-acetylation of H3 provided no affinity enhancement over mono-acetylated H3K14ac (**Table 1**). This is notable because H3 acetylation is the mark most directly associated with transcriptional activation[31, 44, 45]. Our data therefore suggest that the CECR2-BRD acts primarily as a reader of acetylation density on the H4 tail.

Weaker recognition of acetylated H3 may still contribute to chromatin engagement in the context of the nucleosome, where both multi-acetylated H3 and H4 nucleosomes were robustly bound (**Figure 4**), or within the full-length CERF complex. Together, these findings support a role for CECR2 in recognizing hyperacetylated, transcriptionally permissive chromatin[4, 43]. NMR chemical shift perturbation (CSP) analysis provided residue-level insights into the interaction between the CECR2-BRD and multi-acetylated histone H3 and H4 ligands. Significant perturbations were observed in residues located within the ZA and BC loops and the WPF shelf, consistent with engagement of the canonical acetyllysine-binding pocket[16, 46]. Comparable CSP patterns were detected for acetylated histone H3 and H4 ligands, which indicates they are recognized through a conserved mechanism (**Figure 5**). This behavior is similar to what has been reported for several BET-family bromodomains, such as BRD2, BRD3, and BRD4, which also rely on conserved ZA and BC loop dynamics to accommodate different patterns of acetylation with minimal structural rearrangement[16, 47–49]. Mutational analysis further supports a ligand-dependent role for residues within the canonical CECR2 bromodomain binding pocket. Substitution of the ZA-loop aspartate (D464) to alanine enhances the affinity with acetylated H3 ligands, while it reduces the affinity with acetylated H4 ligands. A similar substitution in the conserved asparagine (N514), which typically anchors the acetyllysine carbonyl through a critical hydrogen-bonding interaction[16], had a relatively modest effect on the interaction with acetylated H3, but substantially impaired recognition of both mono- and tetra-acetylated H4 ligands. Similar effects are observed upon mutation of the conserved asparagine residue in other bromodomains such as BRD2-BD1, BRD4-BD1, and BRDT-BD1, where disruption of this residue severely compromises acetyllysine coordination[48–50]. Our results indicate that D464 and N514 contribute to acetylated histone recognition by the CECR2-BRD in a ligand-dependent manner, with N514 playing a particularly important role in H4 recognition.

Next, we characterized the ability of the CECR2-BRD to bind an acetylated version of the non-histone protein RelA. Acetylation of K310 in RelA is required to fully activate the transcriptional functions of NF-κB[9, 51]. CECR2 is a critical epigenetic regulator that interacts with acetylated RelA, and increases the expression of NF-κB target pro-inflammatory genes to drive breast cancer metastasis[10]. Our ITC data confirmed that the CECR2-BRD binds to the mono-acetylated RelA_[305-315]_K310ac ligand, albeit with a weaker affinity than the multiply acetylated histone H3 and H4 ligands (**Tables 1 and 3**). NMR perturbation analysis further demonstrated that there are notable differences in the coordination of mono-acetylated RelA, particularly within residues in the ZA and BC loop regions of the CECR2-BRD, where these unstructured loop regions display larger structural rearrangements upon RelA_[305-315]_K310ac ligand binding (**Figure 6**) compared to histone ligands. This subtle difference in ligand coordination is likely important for the distinction of histone versus non-histone proteins recognized by the CECR2-BRD. The sequence motifs found in non-histone ligands are not as conserved as they are in histone ligands, and the CECR2-BRD must adapt to accommodate recognition of acetylated RelA through additional contacts in the acetyllysine binding pocket to stabilize the interaction.

Huang *et al*. found that the bromodomain of BRD4 also binds to acetylated lysine 310 on RelA, which enhances transcriptional activation of NF-κB[52]. This result suggests overlapping roles between BRD4 and CECR2. However, they did not examine the specific binding mechanism, and it is unclear if BRD4 and CECR2 utilize a conserved binding mode to interact with RelA. Overall, the CECR2-BRD can be placed in the category of bromodomains with broad acetylation selectivity and minimal sequence discrimination. This acetyllysine recognition mode is distinct from the BET family members (e.g., BRD2/4), which exhibit preferential recognition of defined Kac- XX-Kac motifs[53]. While the CECR2-BRD discriminates between histone and non-histone ligands (e.g., RelA), its histone binding profile is consistent with recognition of general acetylation density, especially on the H4 tail, rather than strict sequence motifs.

There are currently two known small molecule inhibitors of the CECR2 bromodomain, namely NVS-CECR2-1 (Structural Genomics Consortium, https://www.thesgc.org/chemical-probes/NVS-CECR2-1) and GNE-886 (Genentech, USA)[54, 55]. Of the two, NVS-CECR2-1 has the most potential because it was demonstrated to inhibit the CECR2-RelA interaction *in vitro*[10], and it blocks the CECR2 bromodomain chromatin binding activity[54]. To begin exploring the structural basis for inhibitor recognition, we attempted NMR titrations with NVS-CECR2-1.

However, interpretation was limited by the requirement of high concentrations of DMSO to solubilize the compound. DMSO alone induced substantial chemical shift perturbations in the CECR2-BRD, particularly within the acetyllysine-binding pocket (ZA and BC loops, **Supp. Figure S11**). This is consistent with prior reports that show DMSO can non-specifically interact with bromodomains[35]. These findings highlight the need for caution when using DMSO-solubilized compounds in BRD-binding assays and reinforce the importance of orthogonal or DMSO-free approaches for mechanistic characterization.

Our results demonstrate the histone-binding preferences of the CECR2-BRD and identify critical residues within its binding pocket that drive the distinct binding modalities observed between histone and non-histone ligands. These results support a model where the CECR2-BRD plays an important role in facilitating the interaction of CECR2 with chromatin where it functions as an epigenetic regulator of gene expression.

However, characterizing the BRD of CECR2 in isolation, rather than the full-length protein, introduces some limitations to the conclusions of our studies. The regulation of gene expression can be extremely nuanced and context-dependent, and while our results inform initial interaction mechanisms between CECR2 and other regulatory proteins, these interactions may differ in the context of the full-length protein. Although ITC revealed a clear preference of the untagged CECR2-BRD for the tetra-acetylated H4 peptide (**Table 1**), the GST-CECR2-BRD bound tetra-acetylated H3 and H4 nucleosomes comparably. This may reflect multivalent binding contributed by the GST tag and bead-based detection in the Captify-Alpha assay, as well as differences in histone tail accessibility within the nucleosome[30–32]. Furthermore, while the use of modified histone peptide ligands is an accepted technique for structural and functional studies, it has additional limitations. These ligands cannot capture the entirety of the interactions (steric, ionic, hydrophobic, etc.) that occur within the context of the nucleosome. Despite these limitations, our results establish a foundation for further in-depth investigations into the role of CECR2 as a regulator of gene expression and inflammation.

The functional implications of non-histone protein acetylation have been explored in various cellular processes[56]. Non-histone protein acetylation regulates p53- mediated apoptosis[57], NF-κB signaling[7], and metabolic reprogramming in cancer cells[56]. In breast cancer, dysregulation of NF-κB signaling drives metastasis[58].

CECR2 was recently recognized as a ‘top epigenetic regulator’ that upregulates NF-κB inflammatory and pro-metastatic genes, emphasizing the therapeutic potential of targeting this bromodomain-containing protein in cancer treatment. Understanding the intricate differences in bromodomain recognition of histone versus non-histone acetylation marks is critical for elucidating the role of these epigenetic mechanisms in the initiation and progression of various diseases. These findings provide strong support for further investigation into CECR2-BRD as a therapeutic target and continued exploration of bromodomain roles in disease contexts.

## CONCLUSION

In conclusion, the results presented in this study shed light on the recognition and specificity determinants of the CECR2-BRD towards acetylation on histone versus non-histone proteins. Our findings demonstrate that the CECR2-BRD exhibits an ability to recognize a broad range of acylated lysine modifications, accommodating diverse combinations of PTMs on histone H3 and H4 tails. Through NMR titration experiments, we found that the CECR2-BRD uses a distinct binding mode to interact with acetylated histones in comparison with an acetylated RelA peptide. Moreover, site-directed mutagenesis experiments identified key residues within the ZA and BC loops that govern the selective recognition of histone versus non-histone substrates. Overall, this study enhances our knowledge of CECR2-BRD function in coordinating specific PTMs, laying the groundwork for further investigations into the molecular mechanisms underlying the bromodomain-mediated epigenetic regulation and its potential implications in disease pathogenesis and therapeutic development.

## AUTHOR CONTRIBUTIONS

M.P. and K.C.G. conceived and designed the experiments. L.F.K. and M.R.M. performed the Captify-Alpha array and analyzed the data. M.P. and E.D.C. expressed and purified the recombinant proteins. M.T. collected the NMR data, and M.P. analyzed the data. B.D. and A.H. collected and analyzed the analytical ultracentrifugation data. M.P., E.D.C., J.M.L., B.D., G.S.S., J.L.S., S.F., and K.C.G. wrote the manuscript, and all authors contributed to editing the manuscript, reviewed the results, and approved the final version.

## FUNDING

Research reported in this study was supported by the National Cancer Institute of the National Institutes of Health under award number (P01CA240685 to K.C.G., G.S.S., J.L.S. and S.F.). This study used the National Magnetic Resonance Facility at Madison, supported by NIH National Institute of General Medical Sciences grant P41GM136463, old numbers P41GM103399 (NIGMS), and P41RR002301. Equipment was purchased with funds from the University of Wisconsin-Madison, the NIH (P41GM103399, S10RR02781, S10RR08438, S10RR023438, S10RR025062, S10RR029220), the NSF (DMB-8415048, OIA-9977486, BIR-9214394), and the USDA. This work was also funded by the Canada 150 Research Chairs program C150-2017-00015, the National Institutes of Health grant 1R01GM120600, and the Canadian Natural Science and Engineering Research Council Discovery Grant DG-RGPIN-2019-05637 (to B.D.). The Canadian Center for Hydrodynamics is funded by the Canada Foundation for Innovation grant CFI-37589 (B.D.). UltraScan supercomputer calculations were supported through NSF/XSEDE grant TG-MCB070039N, and University of Texas grant TG457201 (B.D.). The content is solely the authors’ responsibility and does not necessarily represent the official views of the National Institutes of Health. Automated DNA sequencing was performed in the Vermont Integrative Genomics Resource DNA Facility and was supported by the University of Vermont Cancer Center, the Lake Champlain Cancer Research Organization, and the UVM Larner College of Medicine.

## COMPETING INTERESTS

EpiCypher (M.R.M. and L.F.K.) is a commercial developer of the Captify-Alpha peptide-binding platform used in this study.

## Notes

### Competing Interest Statement

EpiCypher (M.R.M. and L.K.) is a commercial developer of the dCypher peptide-binding platform used in this study.

### Summary of Updates

The revised manuscript was edited in response to reviewer comments.

