## Supplementary material for "The CECR2 bromodomain displays distinct binding modes to select for acetylated histone proteins versus non-histone ligands": Supplmentary information

**Running title**: Acetyllysine Recognition by the CECR2 Bromodomain

**Table of Contents**

Supp. Figure S1: Captify-Alpha analysis of the CECR2 bromodomain…………………...3

Supp. Figure S2: ITC analysis of acetylated histone H3 binding to the CECR2-BRD…..4

Supp. Figure S3: ITC analysis of acetylated histone H4 binding to the CECR2-BRD…..5

Supp. Figure S4: Nucleosome-based Captify assay…………………..……………..……..6

Supp. Figure S5: NMR assignment for the CECR2 bromodomain………………………..7

Supp. Figure S6: NMR titration of CECR2 bromodomain with histone H3K4acK9acK14acK18ac……………………………………………………………………...8

Supp. Figure S7: NMR titration of CECR2 bromodomain with histone H4K5acK8acK12acK16ac……………………………………………………………………...8

Supp. Figure S8: ITC analyses of the interaction between the D464A and N514A mutants of the CECR2 bromodomain and acetylated ligands..…………………….....…...9

Supp. Figure S9: ITC analysis of the binding of RelAK310ac to the CECR2-BRD, D464A, N514A..………………………………………………………….…………………….10

Supp. Figure S10: Interaction of the CECR2 bromodomain with mono-acetylated RelA ligand……………………………………………………………………………………………11

Supp. Figure S11: NMR titration of CECR2 bromodomain with DMSO and NVS-CECR2-1……………………………………………………………………………....…….....11

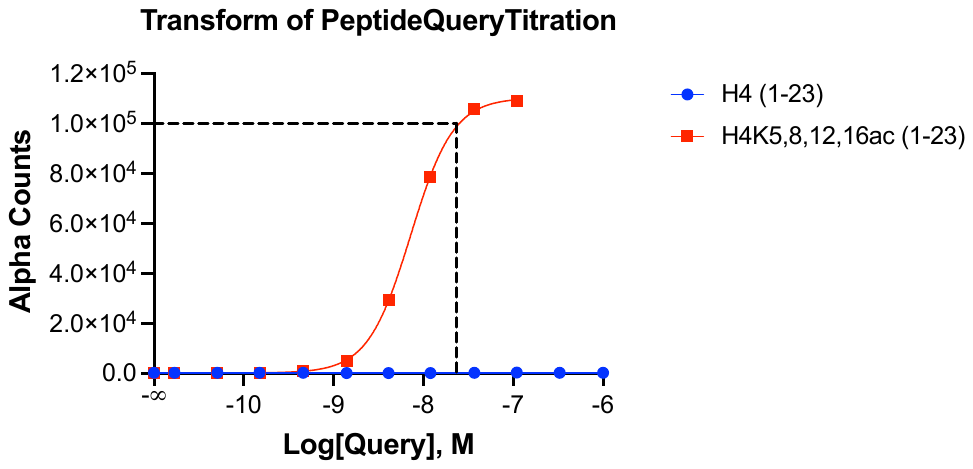

H4_[1-24]_ unmodified

H4_[1-24]_K5acK8acK12acK16ac

**Supp. Figure S1:** Using the Captify-Alpha assay the GST-CECR2-BRD was titrated against unmodified H4_[1-24]_ and H4_[1-24]_K5acK8acK12acK16ac peptides to identify the optimal probing concentration (robust signal-to-background) for peptide panel screens. The dashed line represents the concentration of GST-CECR2-BRD (26 nM) chosen for peptide panel screens (**Figure 1 & Figure 2**).

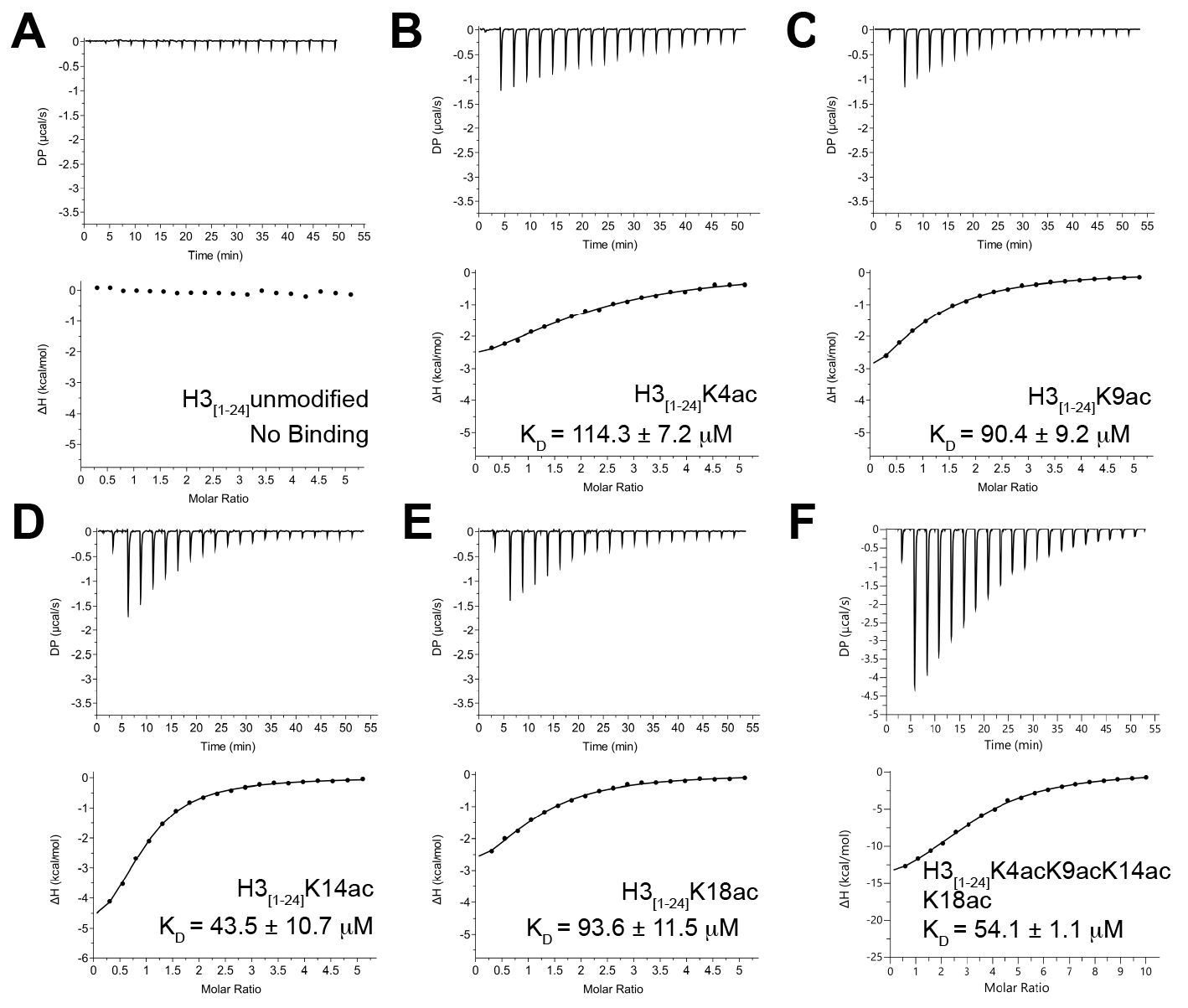

**Supp. Figure S2: ITC analysis of the CECR2 bromodomain binding to acetylated histone H3 peptides.** A-F) Representative ITC titrations of unmodified and acetylated histone H3 ligands into the CECR2 bromodomain. In each panel, the upper trace shows the raw differential power (DP) recorded during sequential injections of peptide into protein, and the lower plot shows the exothermic enthalpy integrated for each injection and plotted against the peptide-to-protein molar ratio. Solid lines represent fits to the integrated binding data. No measurable binding was detected for the unmodified H3 peptide. The apparent dissociation constants (K_D_) shown in each panel are reported as the mean ± standard deviation from at least three independent titrations. All peptides correspond to residues 1-24 of histone H3, and the peptide sequences are provided in Table 1.

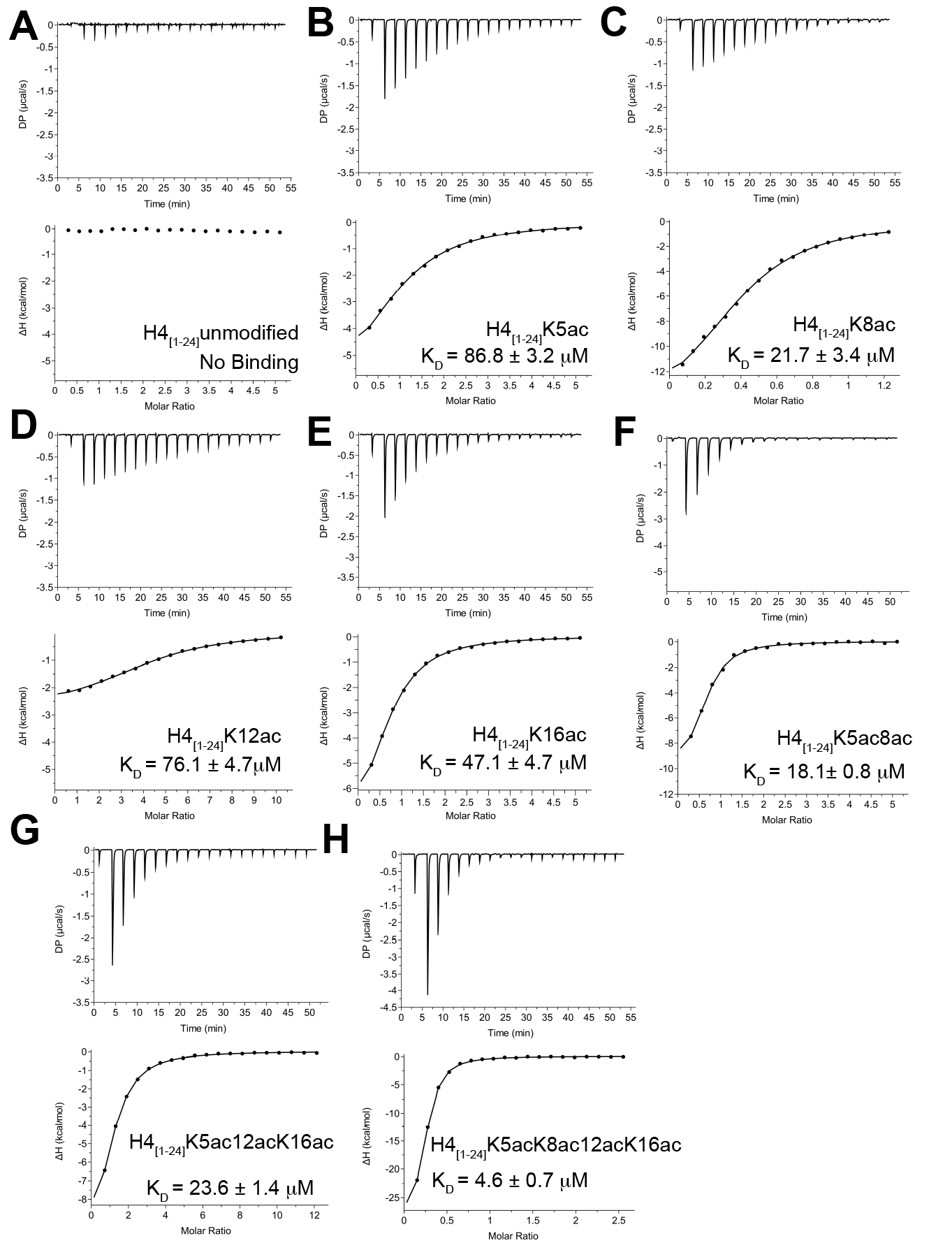

**Supp. Figure S3: ITC analyses of the interaction between the CECR2 bromodomain and acetylated histone H4 peptides.** A-H) Representative ITC titrations of unmodified and acetylated histone H4 ligands into the CECR2 bromodomain. In each panel, the upper trace shows the raw differential power (DP) recorded during sequential injections of peptide into protein, and the lower plot shows the exothermic enthalpy integrated for each injection and plotted against the peptide-to-protein molar ratio. Solid lines represent fits to the integrated binding data. No measurable binding was detected for the unmodified H4 peptide. The apparent dissociation constants (K_D_) shown in each panel are reported as the mean ± standard deviation from at least three independent titrations. All peptides correspond to residues 1-24 of histone H4, and the peptide sequences are provided in Table 1.

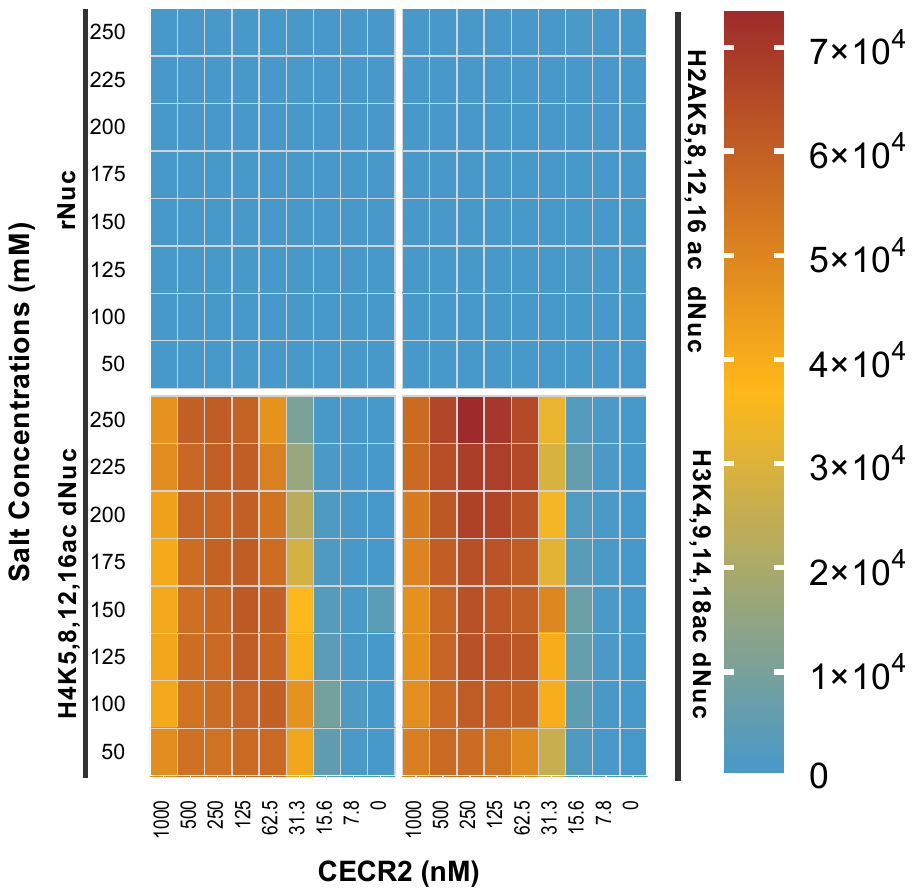

**Supp. Figure S4:** Using a nucleosome-based Captify assay both GST-CECR2-BRD and salt condition (NaCl, mM) were titrated to determine the optimal concentration range and buffer condition. The X-axis represent concentration of GST-CECR2-BRD and Y-axis is concentration of NaCl. Alpha counts are represented by color key.

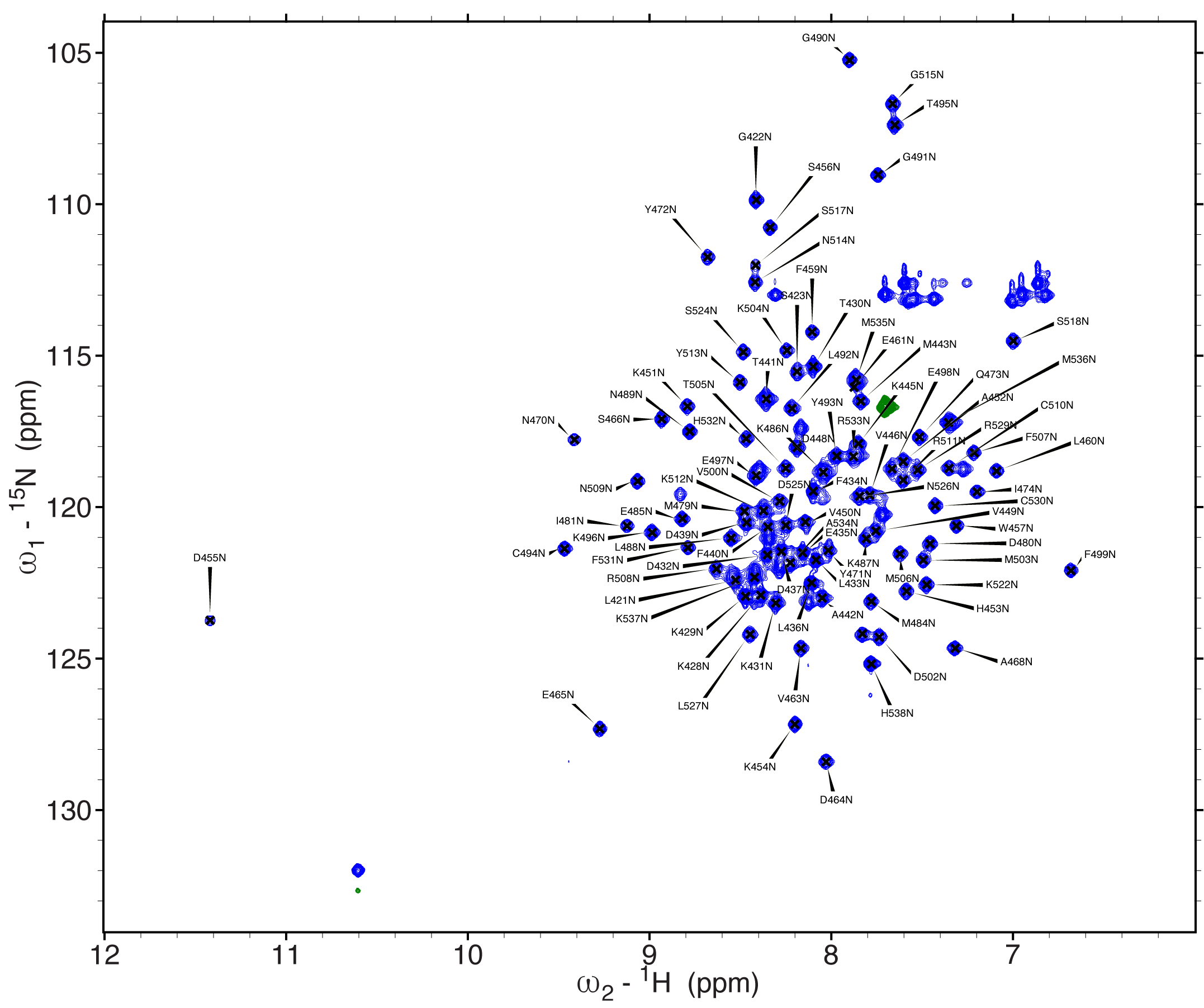

**Supp. Figure S5: 2D ^1^H-^15^N HSQC Spectrum of the CECR2 Bromodomain with assignment.** A total of 106 peaks were observed out of the expected 116, with an assignment rate of approximately 88% (94/116).

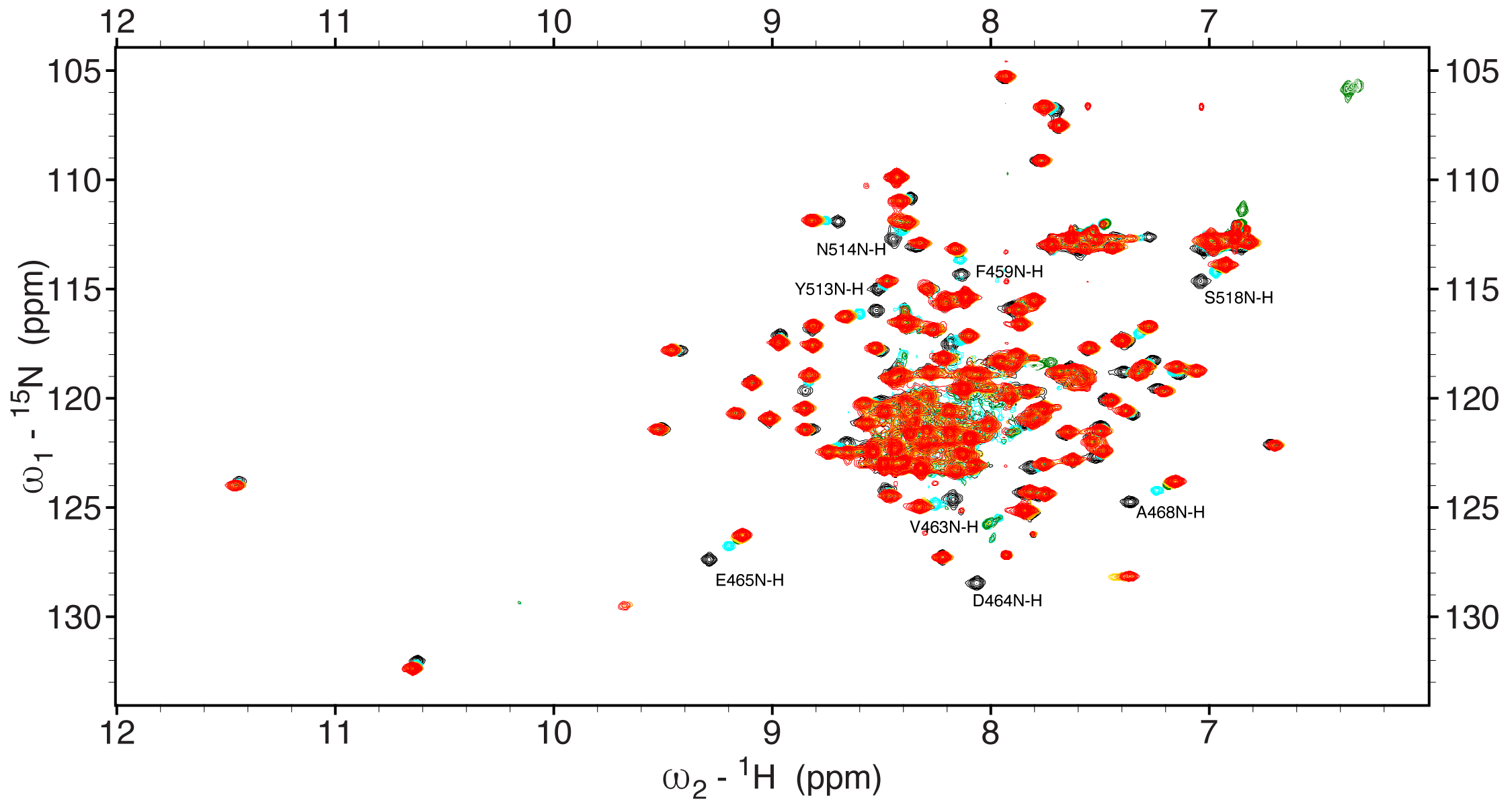

**Supp. Figure S6: Interaction of the CECR2 bromodomain with tetra-acetylated histone H3 ligand.** Superimposed 1H-15N HSQC spectra of the CECR2 bromodomain during titration with the H3K4acK9acK14acK18ac (1-24) peptide at 25°C, with color-coded representations indicating increasing molar concentrations of the peptide - black for Apo CECR2 bromodomain, cyan for 1:0.5, green for 1:1, yellow for 1:2.5, orange for 1:5, and red for 1:10 molar ratio of CECR2 and peptide. CECR2 residues with maximum CSP are labelled.

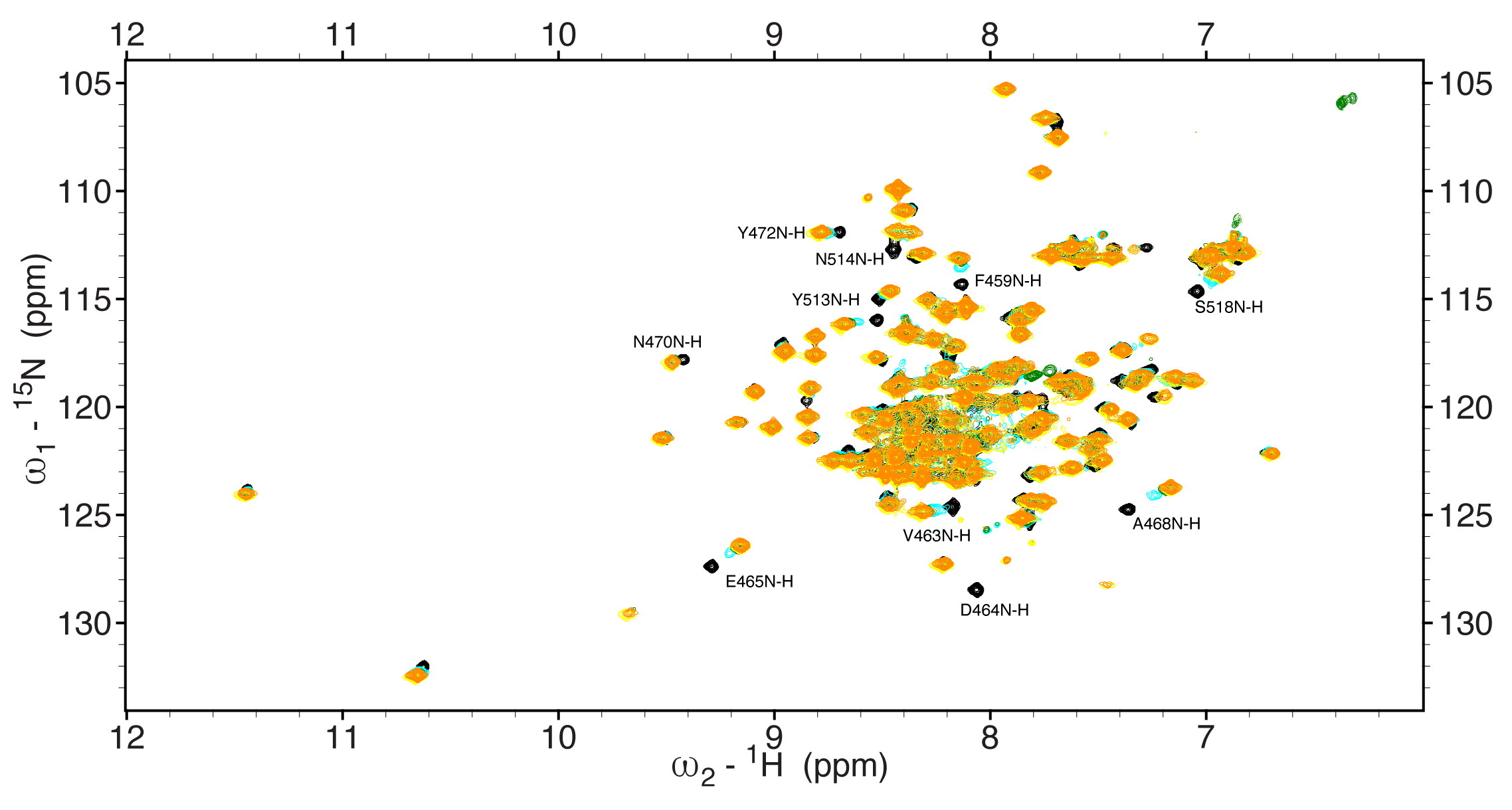

**Supp. Figure S7: Interaction of the CECR2 bromodomain with tetra-acetylated histone H4 ligand.** Superimposed 1H-15N HSQC spectra of the CECR2 bromodomain during titration with the H4K5acK8acK12acK16ac (1-24) peptide at 25°C, with color-coded representations indicating increasing molar concentrations of the peptide - black for Apo CECR2 bromodomain, cyan for 1:0.5, green for 1:1, yellow for 1:2.5, and orange for 1:5 molar ratio of CECR2 and peptide. CECR2 residues with maximum CSP are labelled.

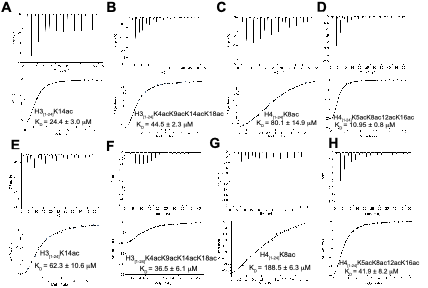

**Supp. Figure S8: ITC analyses of acetylated histone ligand binding to the D464A and N514A mutants of the CECR2 bromodomain.** A-D) Representative ITC titrations of acetylated peptides into the CECR2 D464A bromodomain. E-H) Representative ITC titrations of acetylated peptides into the CECR2 N514A bromodomain. In each panel, the upper trace shows the raw differential power (DP) recorded during sequential injections of peptide into protein, and the lower plot shows the exothermic enthalpy integrated for each injection and plotted against the peptide-to-protein molar ratio. Solid lines represent fits to the integrated binding data. The apparent dissociation constants (K_D_) shown in each panel are reported as the mean ± standard deviation from at least three independent titrations. All peptides correspond to residues 1-24 of the indicated histone, and the peptide sequences are provided in Table 2.

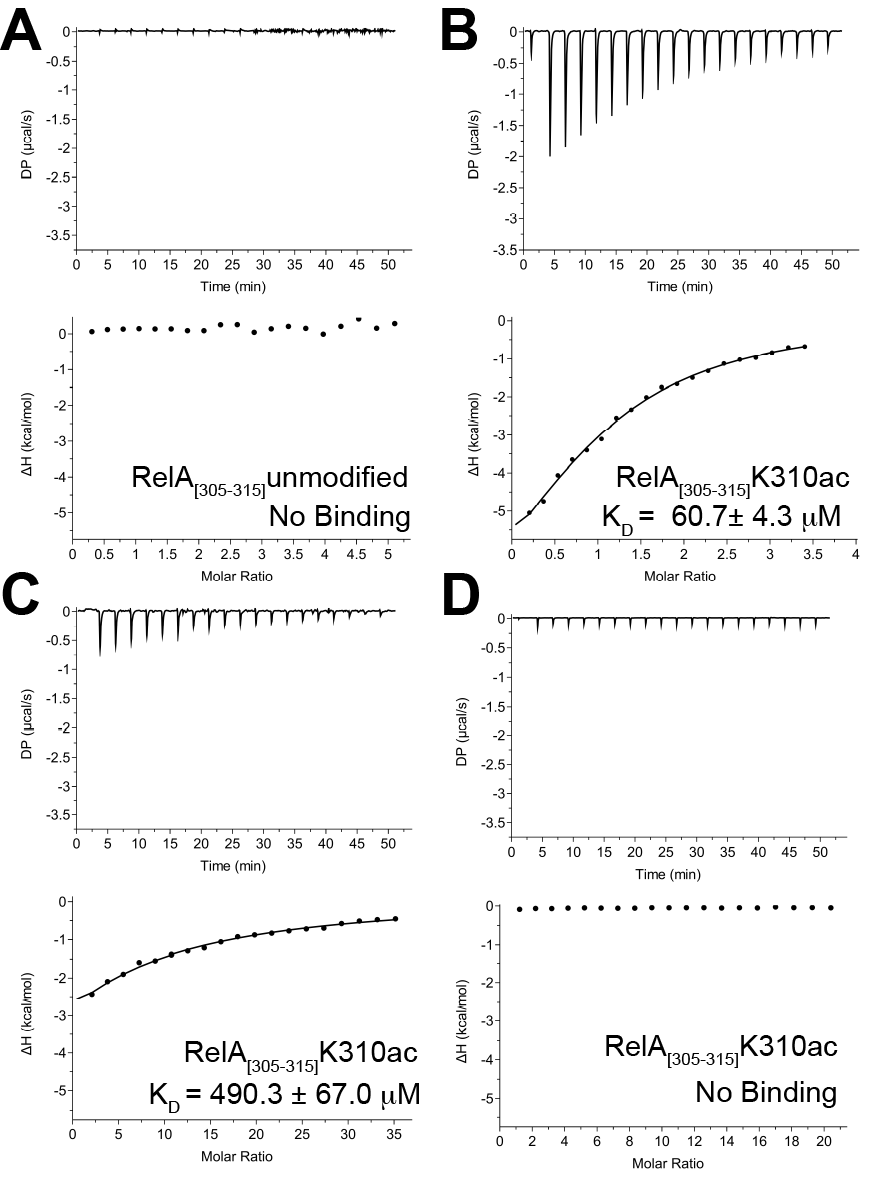

**Supp. Figure S9: ITC analyses of RelAK310ac peptide binding to the wild-type, D464A, and N514A CECR2 bromodomains.** A-B) Representative ITC titrations of the unmodified and K310ac RelA peptide into the WT CECR2 bromodomain. C-D) Representative ITC titrations of the RelA K310ac peptide into the D464A and the N514A CECR2 bromodomains, respectively. In each panel, the upper trace shows the raw differential power (DP) recorded during sequential injections of peptide into protein, and the lower plot shows the exothermic enthalpy integrated for each injection and plotted against the peptide-to-protein molar ratio. Solid lines represent fits to the integrated binding data. No measurable binding was detected for the unmodified RelA peptide. The apparent dissociation constants (K_D_) shown in each panel are reported as the mean ± standard deviation from at least three independent titrations. All peptides correspond to residues 305-315 of RelA, and the peptide sequences are provided in Table 3.

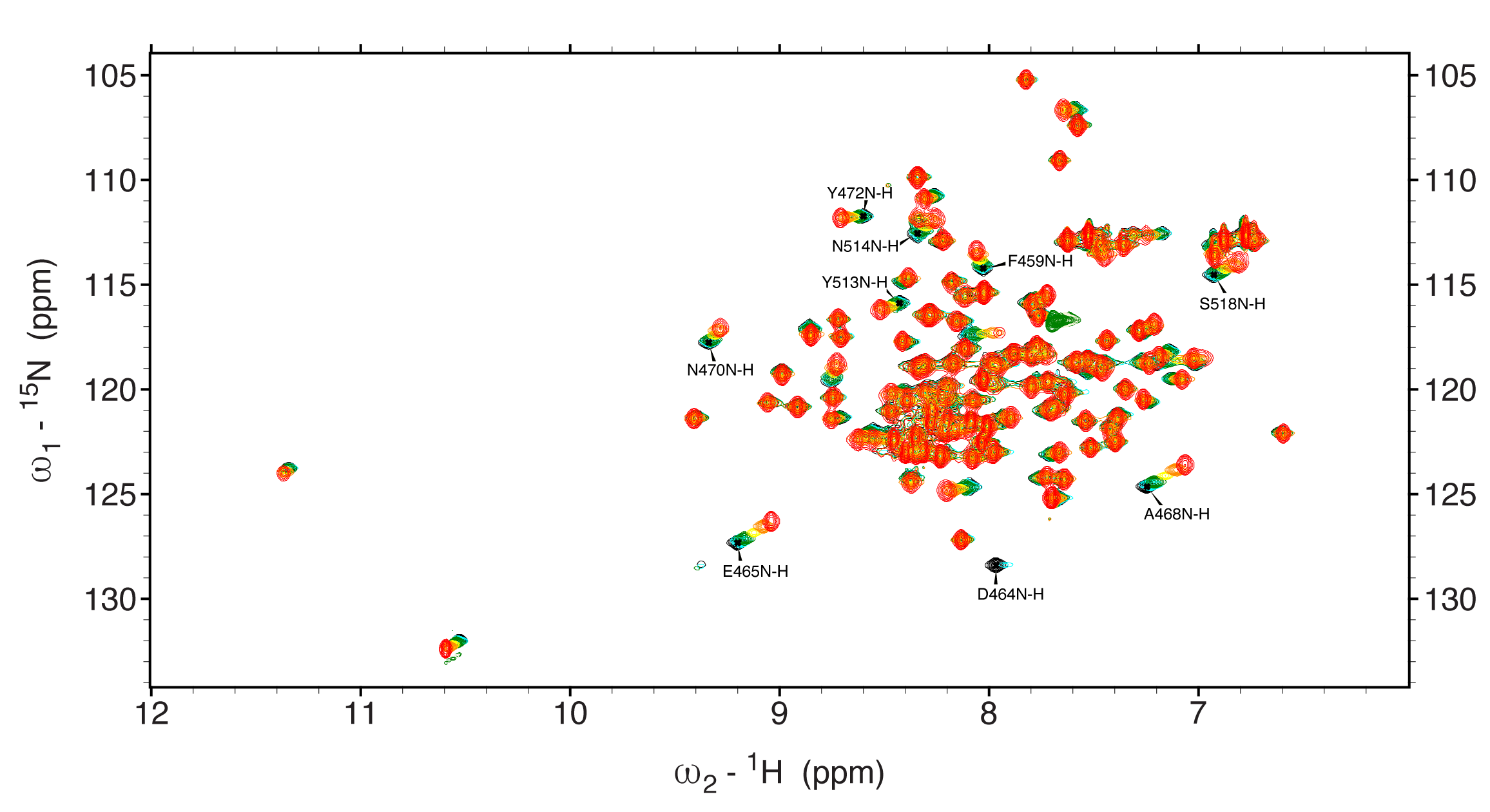

**Supp. Figure S10: Interaction of the CECR2 bromodomain with mono-acetylated RelA ligand.** Superimposed 1H-15N HSQC spectra of the CECR2 bromodomain during titration with the RelAK310ac peptide at 25°C, with color-coded representations indicating increasing molar concentrations of the peptide - black for Apo CECR2 bromodomain, cyan for 1:0.5, green for 1:1, yellow for 1:2.5, orange for 1:5, and red for 1:10 molar ratio of CECR2 and peptide. CECR2 residues with the maximum CSP are labelled.

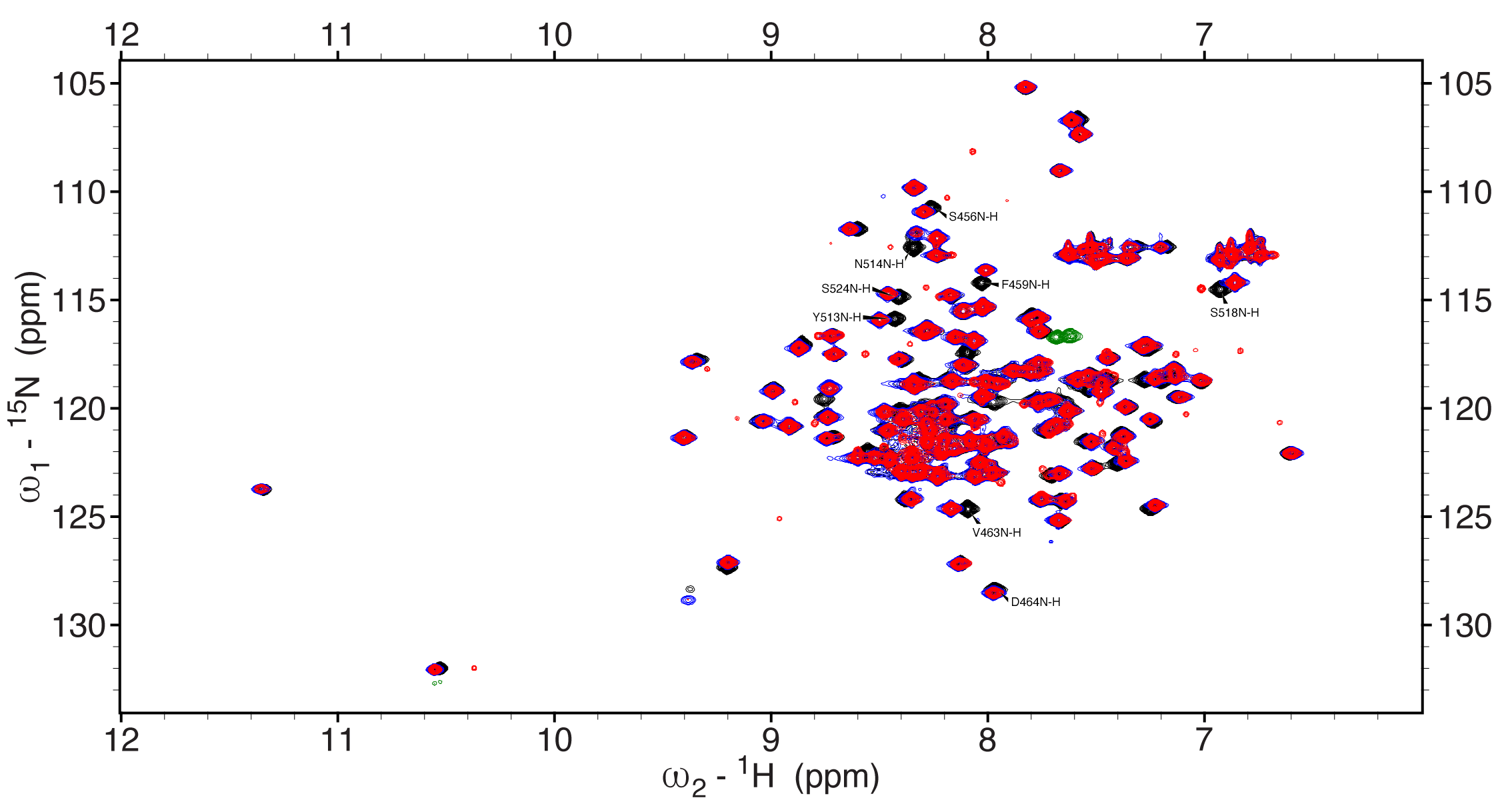

**Supp. Figure S11: Interaction of the CECR2 bromodomain with the CECR2 inhibitor (NVS-CECR2-1). A)** Overlay of the 2D, ^1^H-^15^N HSQC spectra of the CECR2 bromodomain in the absence (black), in the presence of 3% DMSO alone (blue) and upon addition of NVS-CECR2-1 (red) at a 1:1 molar ratio of protein to ligand. All CECR2 residues with maximum CSP are labelled.
